# DNA-dependent binding of nargenicin to DnaE1 inhibits replication in *Mycobacterium tuberculosis*

**DOI:** 10.1101/2021.10.27.466036

**Authors:** Melissa D. Chengalroyen, Mandy K. Mason, Alessandro Borsellini, Raffaella Tassoni, Garth L. Abrahams, Sasha Lynch, Yong-Mo Ahn, Jon Ambler, Katherine Young, Brendan M. Crowley, David B. Olsen, Digby F. Warner, Clifton E. Barry, Helena I.M. Boshoff, Meindert H. Lamers, Valerie Mizrahi

## Abstract

Natural products provide a rich source of potential antimicrobials for use in treating infectious diseases for which drug resistance has emerged. Foremost among these is tuberculosis. Assessment of the antimycobacterial activity of nargenicin, a natural product that targets the replicative DNA polymerase of *Staphylococcus aureus*, revealed that it is a bactericidal genotoxin that induces a DNA damage response in *Mycobacterium tuberculosis* (*Mtb*) and inhibits growth by blocking the replicative DNA polymerase, DnaE1. Cryo-electron microscopy revealed that binding of nargenicin to *Mtb* DnaE1 requires the DNA substrate such that nargenicin is wedged between the terminal base pair and the polymerase and occupies the position of both the incoming nucleotide and templating base. Comparative analysis across three bacterial species suggests that the activity of nargenicin is partly attributable to the DNA binding affinity of the replicative polymerase. This work has laid the foundation for target-led drug discovery efforts focused on *Mtb* DnaE1.

## INTRODUCTION

Claiming an estimated 1.5 million lives in 2020, tuberculosis (TB) remains one of the leading causes of death globally from an infectious disease (WHO, 2021). The severe disruptions to health services wrought by the COVID-19 pandemic are predicted to worsen this grim toll by a further 1 million TB deaths per annum over the next four years (WHO, 2021). In the absence of a highly efficacious vaccine, prolonged chemotherapy with combinations of anti-TB drugs forms the cornerstone of TB control. However, the rise of drug resistance through ongoing evolution and spread of drug-resistant strains of the aetiologic agent, *Mycobacterium tuberculosis* (*Mtb*), is undermining current efforts. This problem, exacerbated by additional treatment delays caused by the pandemic, underscores the urgent need for new TB drugs with distinct mechanisms of action for inclusion in shorter, safer, and more effective drug regimens. The TB drug discovery and development pipeline established in recent years has begun to deliver new and repurposed drugs and combinations which have revolutionized the treatment of drug-resistant TB (Conradie et al., 2020), and demonstrated that treatment-shortening is an achievable goal (Dorman et al., 2021). However, maintaining this momentum requires replenishment of the pipeline with high-quality hit compounds that show mechanistic novelty (Evans and Mizrahi, 2018). This is a key objective of the Tuberculosis Drug Accelerator (TBDA) (Aldridge et al., 2021).

Of the vital cellular processes targeted by TB drugs in clinical use, DNA replication stands out as relatively under-represented (de Wet et al., 2019; Ditse et al., 2017; Reiche et al., 2017); this is despite the high vulnerability of some genes essential for DNA replication in *Mtb* (Bosch et al., 2021) including those encoding DNA gyrase, the target of the fluoroquinolones, moxifloxacin, gatifloxacin and levofloxacin, and the only DNA metabolic enzyme currently targeted for TB therapy. Fluoroquinolones inhibit DNA gyrase with bactericidal consequences for *Mtb* (Mayer and Takiff, 2014; Nagaraja et al., 2017) and have been incorporated in second-line therapy for multidrug resistant (MDR) TB (Dawson et al., 2015) and in treatment-shortening regimens for drug-susceptible TB (Dorman *et al*., 2021). The identification of novel scaffolds that target DNA gyrase remains an active area of investigation (Das et al., 2019; Johnson et al., 2019) while topisomerase I is also being pursued as a new TB drug target (Godbole et al., 2015). Recently, the replisome – the macromolecular machine that copies the bacterial chromosome – has emerged as an attractive potential target for TB (Ditse *et al*., 2017; Reiche *et al*., 2017) and antibacterial drug discovery, more generally (Kaguni, 2018). Key discoveries involving natural products have added impetus to exploring this target further: firstly, griselimycin, a cyclic depsipeptide discovered more than 50 years ago, was shown to bind with high affinity and selectivity to the β-clamp (DnaN) at the site of interaction with DNA polymerase and other DNA metabolizing enzymes (Kling et al., 2015). During DNA replication, the β-clamp interacts with DnaE1, the replicative DNA polymerase termed variously DnaE, DnaE1, or PolC in different bacteria, greatly enhancing the processivity of the polymerase. Griselimycin interferes with the protein interaction between DnaE1 and the β-clamp affecting the processivity of DNA replication (Kling *et al*., 2015). The mechanistic novelty of griselimycin led to the development of the analogue, cyclohexyl-griselimycin, which has improved potency and stability, and demonstrated comparable efficacy to rifampicin when used in combination with first-line drugs in a mouse infection model (Kling *et al*., 2015). Secondly, studies in *Staphylococcus aureus* identified the replicative DNA polymerase, DnaE, as the target of nargenicin A1 (referred to here as nargenicin) (Painter et al., 2015), which belongs to a class of partially saturated alicyclic polyketides comprising an octalin ring (**Figure 1A**) (Cane and Yang, 1985). Nargenicin is an ether-bridged macrolide antibiotic first isolated from various *Nocardia* species almost three decades ago (Celmer et al., 1980; Pidot and Rizzacasa, 2020). It is a narrow-spectrum antimicrobial (Painter *et al*., 2015) with activity against gram-positive bacteria including methicillin-resistant *S. aureus* and *Micrococcus luteus* (Sohng et al., 2008). The identification of *narR/ngnU* (Dhakal et al., 2020; Pidot and Rizzacasa, 2020), a *dnaE* homologue immediately adjacent to the nargenicin biosynthetic gene cluster in the producer organism, *Nocardia* sp. CS682 (Dhakal et al., 2019), suggested a mechanism of self-resistance to nargenicin using NarR/NgnU as a “decoy” (Pidot and Rizzacasa, 2020).

**Figure 1.**
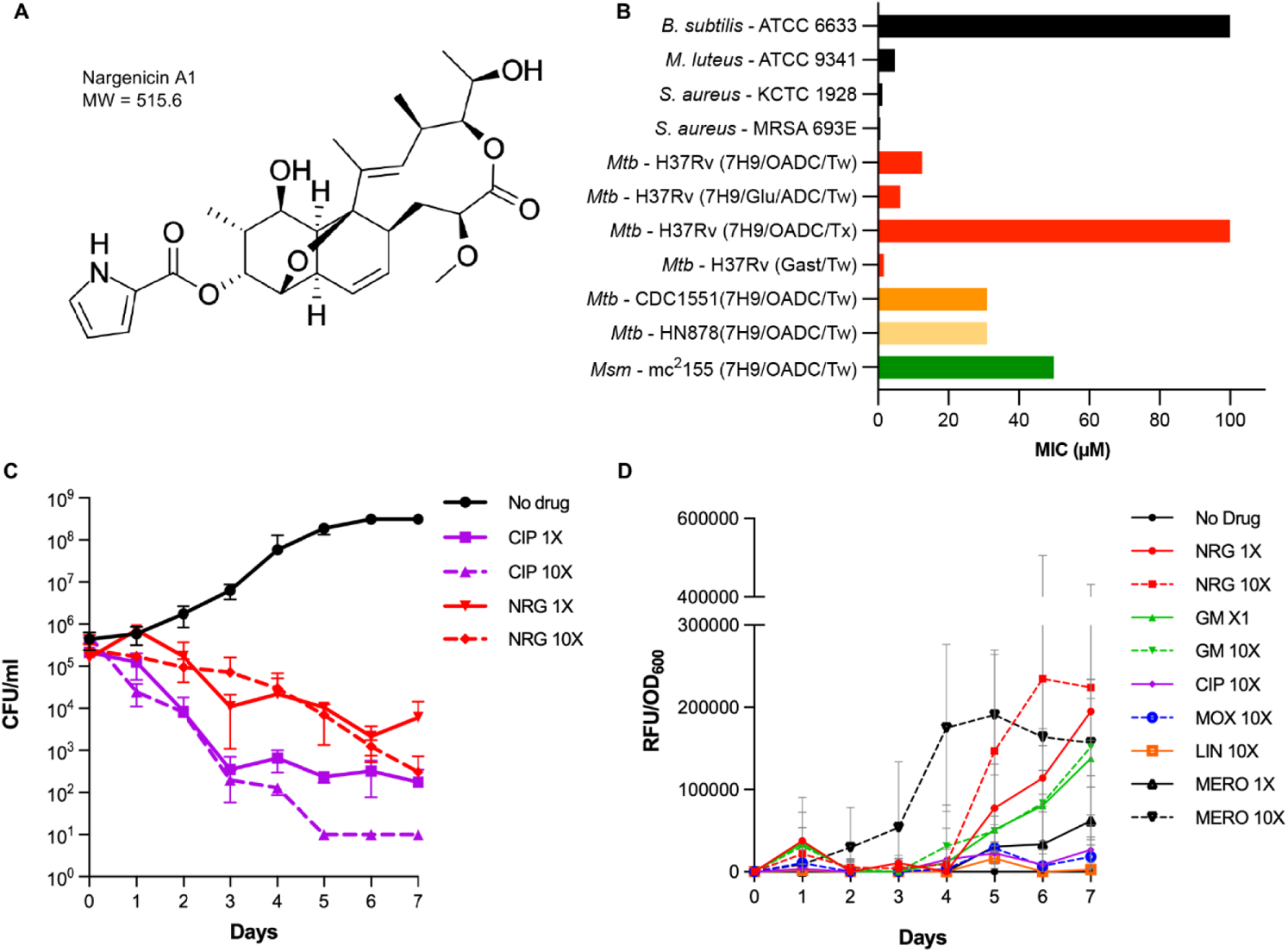
Antimycobacterial activity profile of nargenicin. (A) Chemical structure of nargenicin A1. (B) Antibacterial activity (minimal inhibitory concentration, MIC) of nargenicin (NRG) in mycobacteria and other organisms illustrating the effect of media composition on activity. 7H9, Middlebrook 7H9 media; GAST/(Fe), glycerol alanine salts (with iron); Glu, glucose; (O)ADC, (oleic acid)-albumin-dextrose-catalase; Tw, Tween-80; Tx, Tyloxapol. (C) Time-kill kinetics of nargenicin in *Mtb*, measured by CFU enumeration. Error bars represent the SD derived from two biological replicates. Ciprofloxacin (CIP; MIC = 1.5 µM) was used as a comparator. (D) Drug-induced lytic activity measured by release of GFP from *Mtb* H37Rv-GFP at the indicated concentrations (Chengalroyen *et al*., 2020). Linezolid (LIN; MIC = 1.5 µM) and meropenem (MERO; MIC = 5.2 µM) were used as the non-lytic and lytic controls, respectively. Data are a representative of the two biological replicates.

The potent bactericidal activity and low frequency of resistance for nargenicin in *S. aureus* (Painter *et al*., 2015) led us to investigate the antimycobacterial properties of this molecule (Young et al., 2017) under the auspices of the TBDA. Here, we show that nargenicin is a bactericidal genotoxin that induces a DNA damage response in *Mtb* that is accompanied by cellular elongation and potential weakening of the cell envelope. We further demonstrate that the antimycobacterial activity of nargenicin is mediated through inhibition of DNA synthesis, consistent with inhibition of the DNA polymerase activity of purified DnaE1. Structural analysis by cryo-electron microscopy (cryo-EM) revealed a unique mode of binding by nargenicin to *Mtb* DnaE1 in the presence of DNA in which nargenicin occupies the position of both the incoming nucleotide and templating base and stacks onto the terminal base pair. We show that the antibacterial efficacy of nargenicin as a DNA replication inhibitor is attributable, at least in part, to the DNA binding affinity of the organism’s replicative polymerase.

## RESULTS

### Nargenicin is bactericidal against *Mtb in vitro*

Nargenicin was shown to have a minimum inhibitory concentration (MIC) of 12.5 µM against *Mtb* H37Rv under standard culture conditions (7H9/OADC/Tw) (**Figure 1B; Table S1**). In this culture medium, nargenicin showed comparable activity against a range of drug-sensitive and drug-resistant clinical isolates of *Mtb* and was active against *M. smegmatis* (*Msm*). The activity against *Mtb* diminished significantly when Tween-80 was replaced by Tyloxapol to disperse the mycobacteria. This likely reflects the differential impact of these two detergents on the lipid composition of the cell envelope at the concentrations typically used for clump dispersal (Ortalo-Magne et al., 1996) with Tween-80 increasing permeability to the drug (Tullius et al., 2019). Nargenicin showed increased potency in GAST/(Fe)/Tween-80. The *in vitro* selectivity index was reasonable with limited cytotoxicity against the HepG2 cell line (CC50 >100 µM).

Time-kill kinetic analysis revealed that nargenicin was bactericidal in *Mtb* H37Rv, showing time-dependent kill with limited dose dependency over the concentration range tested 5 (**Figure 1C**). To ascertain whether this bactericidal activity was accompanied by cell lysis, we quantified GFP release from H37Rv-GFP (Chengalroyen et al., 2020; Kumar et al., 2012). Nargenicin treatment led to GFP release from day 4 onwards, peaking on day 6-7 (**Figure 1D**). Griselimycin treatment also resulted in delayed GFP release analogous to that elicited by nargenicin, but no release of GFP was observed upon exposure to the DNA gyrase inhibitors, ciprofloxacin or moxifloxacin, demonstrating that the GFP release was not a generic consequence of disrupting DNA metabolism (**Figure 1D**).

### Nargenicin inhibits DNA synthesis and is genotoxic in *Mtb*

To ascertain whether nargenicin shares the same mechanism of action in mycobacteria as in *S. aureus* (Painter *et al*., 2015), we applied a suite of complementary biological profiling assays in *Mtb* and *Msm*. Multiple attempts to isolate spontaneous nargenicin-resistant mutants in *Mtb* or *Msm* by plating 10^9^-10^10^ cells on media containing nargenicin at 5-20× MIC (*Mtb*) or 1-10× MIC (*Msm*) were unsuccessful, yielding no heritably resistant mutants. Reasoning that nargenicin would elicit a DNA damage response if it disrupts DNA replication, we used the *Mtb* P*recA*-LUX reporter strain to monitor activity of the DNA-damage-inducible *recA* promoter in response to drug treatment (Naran et al., 2016). Like fluoroquinolones and griselimycin, nargenicin triggered dose-dependent induction of luminescence (**Figures 2A and S1**). Comparative DNA microarray analysis revealed a transcriptomic signature for nargenicin-treated *Mtb* that shared key features with those elicited by mitomycin C and fluoroquinolones (**Figure S2A, S2B and Table S2**) (Boshoff et al., 2004; Boshoff et al., 2003). Genome-wide transcriptome analysis by RNA-seq revealed profound upregulation of *dnaE2*, *imuA*’ and *imuB,* components of the mycobacterial “mutasome” responsible for DNA damage tolerance and SOS-induced mutagenesis (Boshoff *et al*., 2003; Warner et al., 2010), and other DNA repair genes including *recA*, *radA*, *uvrA*, *lhr*, and *adnAB* (**Figures 2B; S2C and Data S1**).

**Figure 2.**
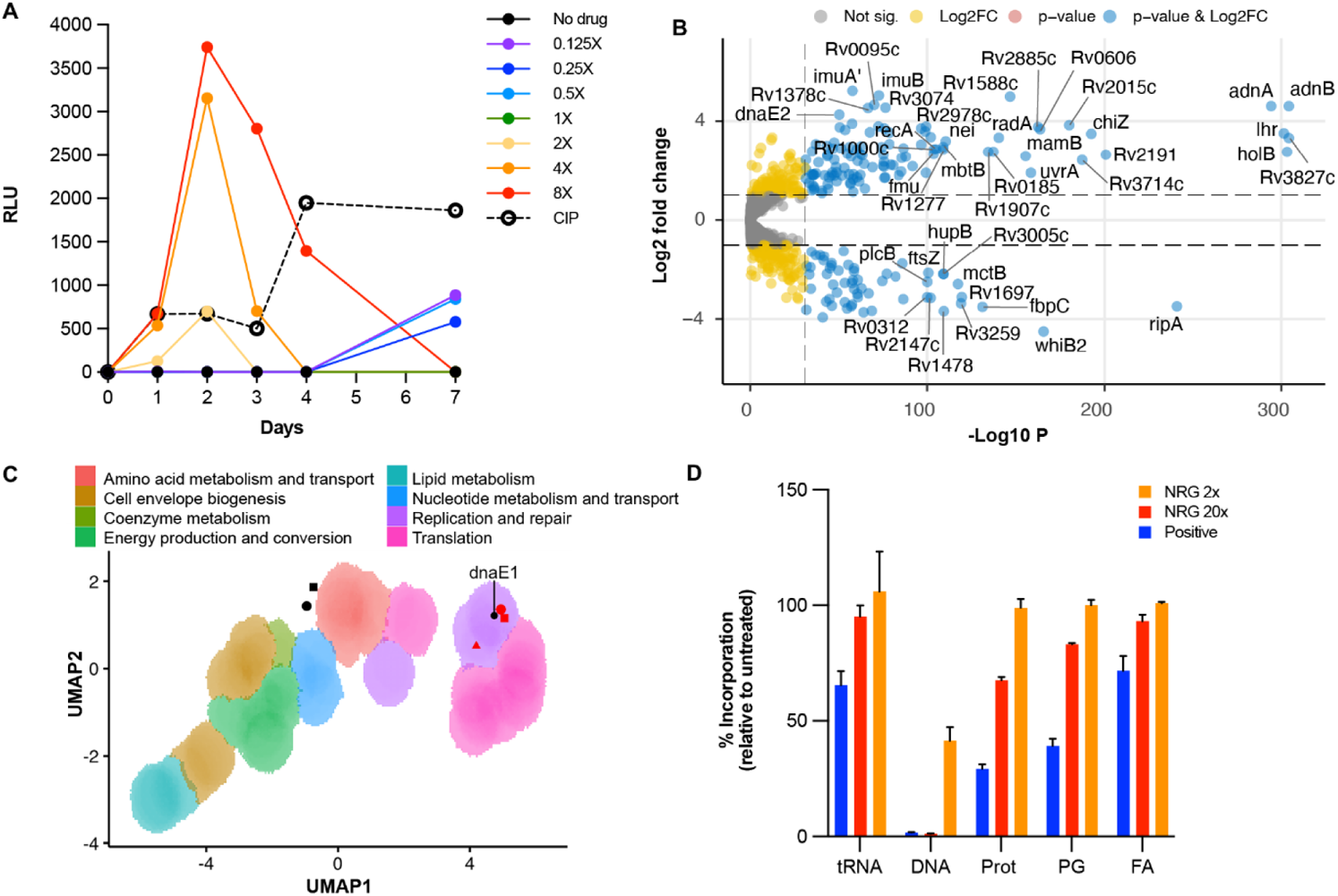
Nargenicin is a genotoxin that inhibits DNA replication in mycobacteria. (A) Analysis of *recA* promoter activity elicited by nargenicin using the reporter strain, PrecA-LUX (Naran *et al*., 2016). Ciprofloxacin (CIP, 2× MIC) was a positive control. RLU, relative luminescence units. (B) Volcano plot illustrating the transcriptional response (RNA-seq) of *Mtb* to nargenicin (10× MIC). Differential expression (Log_2_ fold-change) of nargenicin-treated cultures versus DMSO-treated controls are plotted against adjusted P values (P-value) for each gene indicating significant upregulation of genes involved in the response of *Mtb* to DNA-damaging agents (Boshoff *et al*., 2003). (C) Morphological profiling of *Msm* in response to treatment with nargenicin illustrates that bacillary morphotypes (de Wet *et al*., 2020) cluster in UMAP space with those of CRISPRi hypomorphs in genes involved in DNA replication, including *dnaE1*. Black circle, untreated; black square, DMSO-treated; red circle, nargenicin-treated at 1× MIC; red square, nargenicin-treated at 2× MIC; red triangle, nargenicin-treated 4× MIC. (D) Selective inhibition of DNA synthesis by nargenicin in *Mtb*. The incorporation of radiolabeled precursors into total nucleic acid (tNA), protein (Prot), peptidoglycan (PG), and fatty acid (FA) was measured in the absence (DMSO) or presence of nargenicin at 2× or 20× MIC (black and red bars, respectively). The level of radiolabel incorporation into each macromolecular species is depicted relative the DMSO-treated control. Assay specificity was confirmed using pathway-specific antibiotics as positive controls: ofloxacin (5 μg/mL), streptomycin (10 μg/mL), D-cycloserine (5 μg/mL), and isoniazid (0.2 μg/mL). Error bars represent the standard deviations from two experimental repeats.

Interestingly, deletion of either *recA* (Machowski et al., 2007) or *dnaE2* (Boshoff *et al*., 2003; Warner *et al*., 2010) had a negligible impact on the antimycobacterial activity of nargenicin (**Table S1**). Genes most highly downregulated by nargenicin were enriched in those associated with cell division (*ftsZ*, *whiB2* and *ripA*), and included genes involved in cell envelope biogenesis (*e.g*., *fbpC*) (**Figures 2B and S2C**).

Morphological profiling of *Msm* exposed to nargenicin revealed a filamentation phenotype with the proportion of elongated bacilli in the population increasing with drug dose (**Figure S3**). This drug-induced profile clustered closely in UMAP space with those resulting from transcriptional silencing of components of the DNA replication and repair machinery (**Figure 2C**), as previously defined (de Wet et al., 2020), further implicating disruption of DNA metabolism in the mode of action of nargenicin. Direct evidence for inhibition of DNA replication was then obtained from a macromolecular incorporation assay, which compares incorporation of radiolabeled precursors into total nucleic acid, DNA, protein, peptidoglycan or fatty acid in cells treated with an experimental drug *versus* controls. Nargenicin had a profound effect on DNA synthesis resulting in 60% and >95% reduction in [^3^H]-uracil incorporation when used to treat *Mtb* at 2× and 20× MIC, respectively. In contrast, nargenicin had a limited impact on RNA, protein, peptidoglycan, and fatty acid synthesis (**Figure 2D**). Together, these results were consistent with the replicative polymerase, DnaE1, as the likely target of nargenicin in mycobacteria.

To investigate this further, we assessed the impact of modulating the level of *dnaE1* expression on susceptibility of mycobacteria to nargenicin. We generated a set of fluorescently labeled *Mtb* hypomorphs carrying inducible *dnaE1* CRISPR interference (Rock et al., 2017) constructs and determined the inhibitory activity of nargenicin against these strains in the presence or absence of the anhydrotetracycline (ATc) inducer. Marked hypersensitization to nargenicin was observed for all four hypomorphs under conditions of *dnaE1* silencing (+ATc) but not in the uninduced controls (-ATc) (**Figures 3A-C**). Notably, the effect was specific to nargenicin, as evidenced by the lack of effect of *dnaE1* silencing on the susceptibility of *Mtb* to isoniazid or ciprofloxacin, which target mycolic acid biosynthesis and DNA gyrase, respectively (**Figure 3C**). Together, these results identified DnaE1 as a target of nargenicin in *Mtb*. Interestingly, overexpression of *Msm dnaE1* had no effect on the nargenicin susceptibility in *Msm* or *Mtb* (**Figure S4**) suggesting that DnaE1 copy number alone did not determine nargenicin efficacy.

**Figure 3.**
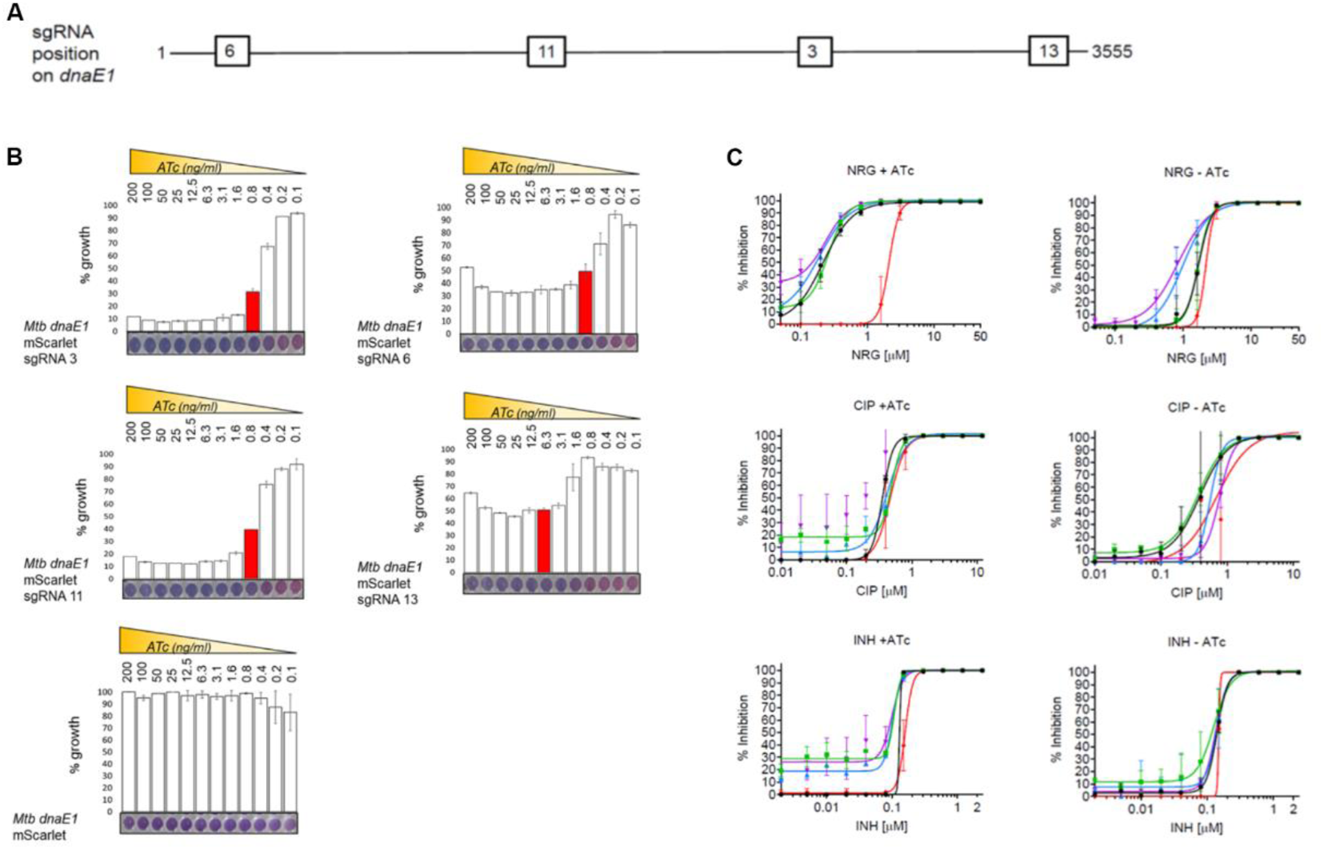
Transcriptional silencing of *dnaE1* by inducible CRISPRi selectively hypersensitizes *Mtb* to nargenicin. (A) Location of sgRNAs 3, 6, 11 and 13 on the *Mtb dnaE1* gene (not drawn to scale). (B) *In vitro* growth phenotypes of the four inducible CRISPRi hypomorphs in *dnaE1* constructed in a strain of *Mtb* carrying a constitutively expressed mScarlet reporter. Strain growth was measured using a microplate alamar blue assay after 7 days’ exposure to ATc at a concentration ranging from 0.1-200 ng/ml. Columns highlighted in red represent the IC50 for ATc. Data plotted represent the average and standard deviation of two technical replicates for one of two independent experiments. (C) The four *dnaE1* hypomorphs were tested for susceptibility to nargenicin (NRG) alongside the control drugs, ciprofloxacin (CIP) and isoniazid (INH). Drug-mediated growth inhibition of the *Mtb dnaE1* mScarlet sgRNA 3 (black), *Mtb dnaE1* mScarlet 6 (green), *Mtb dnaE1* mScarlet sgRNA 11 (blue), *Mtb dnaE1* mScarlet sgRNA 13 (purple) hypomorphs, and *Mtb* mScarlet vector control (red) strains in the presence (+ATc, 100 ng/ml) or absence of inducer (-ATc) was determined by measuring fluorescence intensity at day 14. Data represent the average and standard error of two technical replicates for one representative experiment, fitted with a dose response curve (nonlinear regression model). Experiments were performed in triplicate.

### Nargenicin differentially inhibits bacterial polymerases

Based on the microbiological evidence, we investigated whether nargenicin inhibited the DNA polymerase activity of *Mtb* DnaE1 in a biochemical assay. For comparison, we included *S. aureus* DnaE, as well as the extensively characterized replicative DNA polymerase from *E. coli*, DNA polymerase III α (Pol IIIα). To monitor the polymerase activity, we used a real-time polymerase assay in which the incorporation of dGMPs in the primer strand quenches the fluorescent signal of a fluorescein group at the 5’ end of the template strand (Rock et al., 2015). We found that nargenicin also inhibits the activity of *Mtb* DnaE1, albeit at ~20-fold higher concentrations than *S. aureus* DnaE (IC_50_ = 125 nM and 6 nM, respectively) under the conditions of this assay (**Figure 4A**). Surprisingly, the *E. coli* polymerase was only significantly inhibited by nargenicin at concentrations higher than 10 µM.

**Figure 4.**
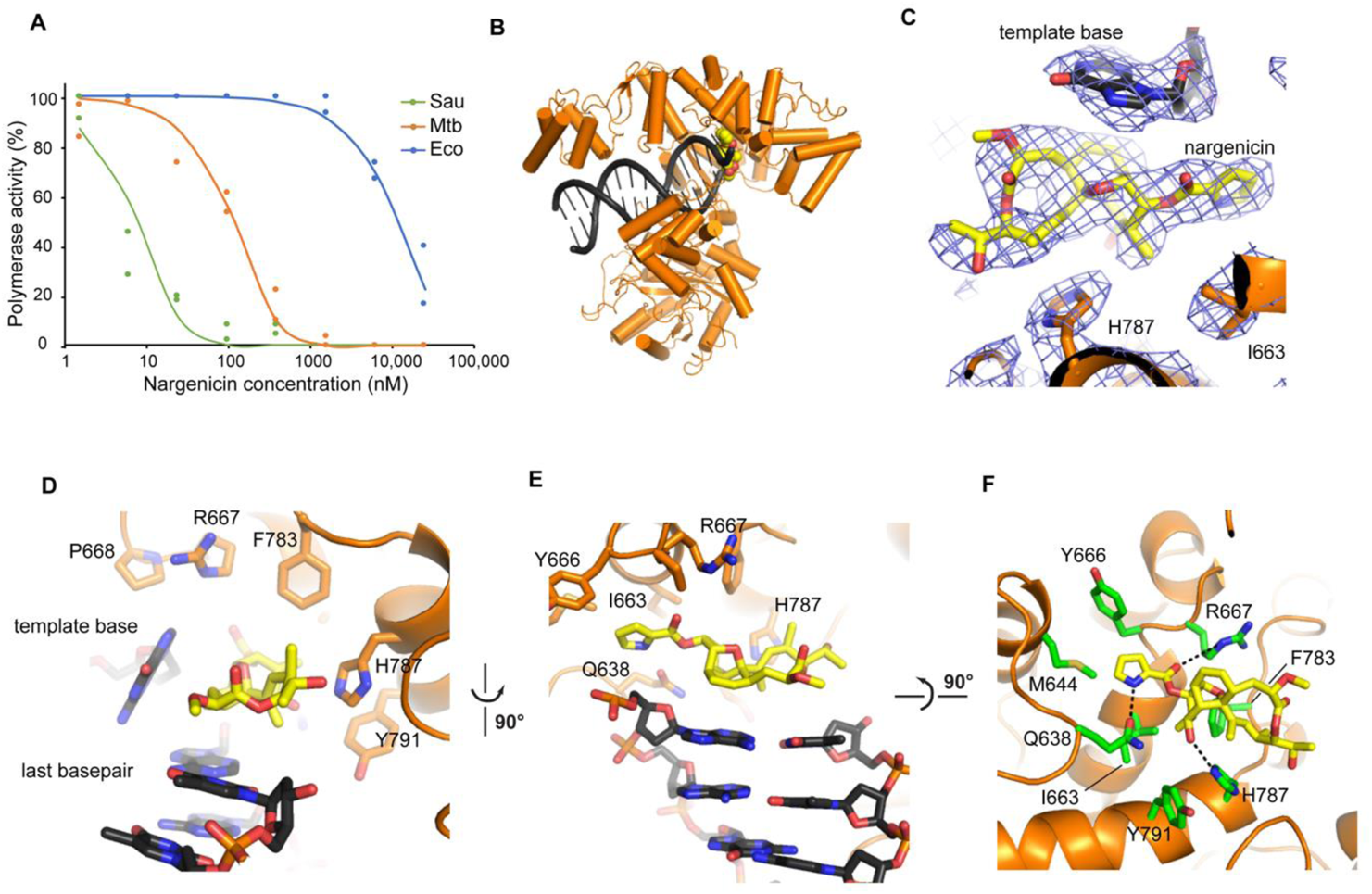
Mechanism of DNA polymerase inhibition by nargenicin. (A) Nargenicin inhibition curves of three bacterial replicative DNA polymerases, *S. aureus* DnaE (green line), *Mtb* DnaE1 (orange line), and *E. coli* Pol IIIα (blue line), show IC50 values of 8 nM, 125 nM, and 13 000 nM, respectively. (B) Cryo-EM structure of *Mtb* DnaE1 bound to DNA and nargenicin in yellow. (C) Magnified view of the nargenicin molecule located between the displaced template base and His787. Cryo-EM map is shown in blue mesh. (D) The composite binding site of nargenicin between the last base pair of the DNA duplex, the displaced templated base, and the fingers domain of the polymerase. (E) Top view of the binding site showing the ‘base paring’ of nargenicin onto the last base pair of the DNA duplex (ssDNA overhang not shown for clarity). (F) The nargenicin binding pocket in DnaE1 as viewed from the DNA. All residues located with 5 Å of nargenicin are shown in green sticks. Hydrogen bonds between the protein and nargenicin are indicated with black dashed lines.

### Cryo-EM reveals mechanism of inhibition by nargenicin

To elucidate the mechanism of polymerase inhibition, we determined the structure of full-length *Mtb* DnaE1 in complex with nargenicin and a DNA substrate by cryo-EM (**Figures 4B-F**). The structure was determined to a resolution of 2.9 Å with well-defined density for the polymerase active site, DNA, and the bound nargenicin molecule (**Figures 4B-F**). The cryo-EM structure of *Mtb* DnaE1 is identical to the previously determined crystal structure (Baños-Mateos et al., 2017) with the exception of the oligonucleotide/oligo saccharide binding (OB) domain that was not included in the crystal structure (**Figure S5**). The OB domain is flexible as it shows a weaker density in the cryo-EM map when compared to the rest of the molecule. The flexibility of the OB domain is consistent with cryo-EM structures of *E. coli* Pol IIIα that show a 70 Å movement of the OB-domain between the DNA-bound and DNA-free state (Fernandez-Leiro et al., 2015).

The DNA is bound in a canonical manner between the thumb and fingers domains, as was previously observed for other C-family DNA polymerases (Evans et al., 2008; Fernandez-Leiro *et al*., 2015; Wing et al., 2008). The nargenicin molecule is bound in the polymerase active site and is sandwiched between the last base pair of the DNA duplex, the first base of the template strand, and the fingers domain of the polymerase (**Figure 4D**). Nargenicin occupies both the position of the incoming nucleotide as well as the template base and thus mimics the position of the newly synthesized base pair (**Figure 4E**). To do so, the first unpaired template base is displaced from its position and bumps into Pro668 of an adjacent helix (residues 668 to 673) that becomes disordered. On the protein side, nargenicin occupies a shallow pocket and only makes three direct contacts with the protein: Arg667 and His787 make a hydrogen bond to two oxygens in nargenicin, while Gln638 makes a hydrogen bond with the nitrogen in the pyrrole ring (**Figure 4F**). The opposite end of nargenicin that is located on top of His787 makes no interaction with the protein as its nearest neighbor is over 5 Å away.

The binding of nargenicin is reminiscent of the binding of aphidicolin in human DNA polymerase α (hPolα) (Brundret et al., 1972). Although the two inhibitors are different in structure (**Figure S6A**) and the polymerases belong to different families (hPolα is a B-family polymerase, whereas *Mtb* DnaE1 a C-family polymerase), both inhibitors are bound between the last base pair of the DNA and the polymerase fingers domain, occupy the position of both incoming and templating base, and displace the templating base (**Figure S6B-C**). However, owing to the structural differences in the polymerase active sites, it is unlikely that nargenicin can inhibit the human polymerase as modeling of nargenicin into the hPolα structure reveals several clashes with the protein (**Figure S6D**. The similar mechanism of action of the two inhibitors derived from different organisms – aphidocolin is derived from the mold, *Cephalosporium aphidicola* (Brundret *et al*., 1972) whereas nargenicin is produced by a *Nocardia* species (Cane and Yang, 1985; Celmer *et al*., 1980) – is a remarkable case of convergent evolution.

### Drug resistance through allostery

The structure described above shows that the DNA forms a crucial part of the nargenicin binding site, agreeing with the previous observation that binding of nargenicin to *S. aureus* DnaE only occurs in the presence of DNA (Painter *et al*., 2015). This DNA dependency of binding may also hold the key to the differences in inhibition between *S. aureus* DnaE, *Mtb* DnaE1, and *E. coli* Pol IIIα (**Figure 5**). The predicted nargenicin binding sites for *S. aureus* DnaE and *E. coli* Pol IIIα are highly similar to those of *Mtb* DnaE1 (**Figures 5A-B**) and the three residues that make a hydrogen bond with nargenicin are conserved in all three species. Hence, the difference in sensitivity does not appear to have its origin in the binding site. Moreover, a mutation in *S. aureus* DnaE (a serine to leucine mutation at position 765, equivalent to *Mtb* DnaE1 residue 860) that renders it resistant to nargenicin is located ~30 Å away from nargenicin (**Figure 5C**). This mutation is immediately adjacent to the region of the fingers domain that interact with phosphate backbone of the double-stranded DNA substrate. Therefore, we hypothesized that the potency of nargenicin to inhibit a DNA polymerase may be dictated by the polymerase’s affinity for DNA. To test this, we measured the DNA affinity of the three polymerases by fluorescence anisotropy using a primed DNA substrate (**Figure 5D**). The three polymerases show strikingly different dissociation constants of ~ 6 nM for *S. aureus* DnaE, 250 nM for *Mtb* DnaE1, and 12 µM for *E. coli* Pol IIIα. These DNA affinities correlate with the relative sensitivities to nargenicin that follow the same trend (**Figure 4A**). We also tested the resistant mutation in *S. aureus* DnaE (S765L), which, as predicted, reduced the affinity for DNA approximately 14-fold (**Figure 5D**).

**Figure 5.**
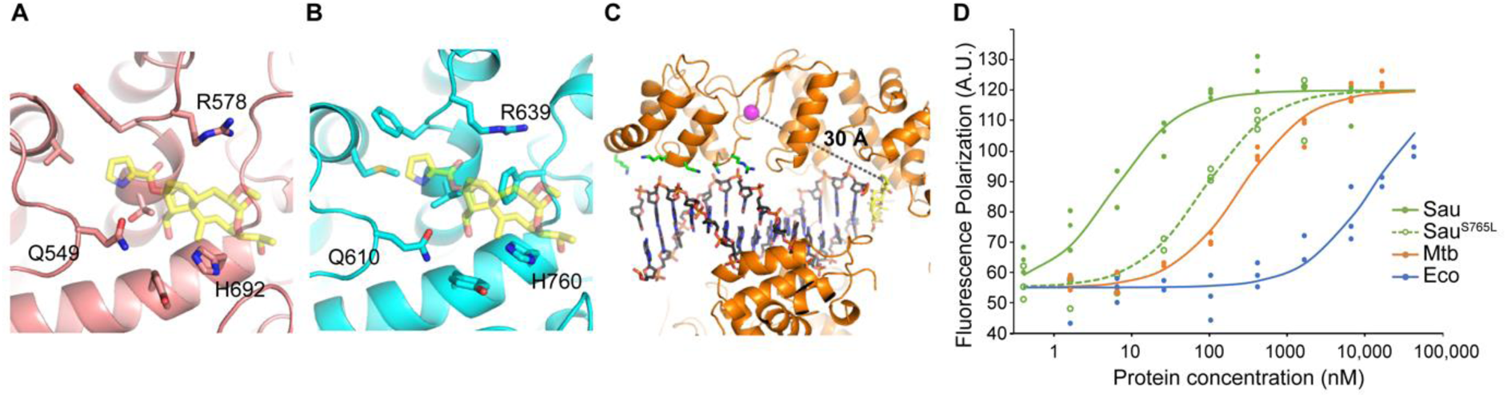
Sensitivity to nargenicin is dependent on DNA binding affinity. (A) Nargenicin binding site in a computational model of *S. aureus* DnaE. (B) Nargenicin binding site in the crystal structure of *E. coli* Pol IIIα. Nargenicin is shown in transparent sticks and the three residues that make a hydrogen bond to nargenicin in *Mtb* DnaE1 are labelled (see also Figure 4F). (C) The nargenicin resistance mutation in *S. aureus* DnaE mapped onto *Mtb* DnaE1, shown by a magenta sphere, is located 30 Å away from the nargenicin (shown in yellow sticks) but is adjacent to the dsDNA binding region of the polymerase. Residues that interact with the DNA backbone are shown in green sticks. (D) Fluorescence anisotropy DNA binding curves of *S. aureus* DnaE (green line), *Mtb* DnaE1 (orange line), and *E. coli* Pol IIIα (blue line) show dissociation constants of 6 nM, 250 nM and 12 µM, respectively. *S. aureus* DnaE^S765L^ (green dashed line) which carries a mutation that confers antibiotic resistance, shows a dissociation constant of 85 nM, which is ~14-fold increased, as compared to wild type.

Taken together, the data support the notion that the potential of nargenicin to inhibit a DNA polymerase is dependent on the polymerase’s affinity for DNA, and any changes which reduce the DNA affinity, lead to reduced nargenicin potency, either through natural variation, as in the case of *E. coli* Pol IIIα, or through a resistance-conferring mutation (Painter *et al*., 2015), as for *S. aureus* DnaE. Importantly, *S. aureus* engages two essential DNA polymerases at the replication fork, namely, PolC and DnaE (Inoue et al., 2001); if the activity of one is impaired, the other may compensate. However, mycobacteria rely on only one replicative polymerase, DnaE1. Therefore, nargenicin resistance-conferring mutations in DnaE1 could have catastrophic consequences in mycobacteria, which might explain our inability to isolate spontaneous resistant mutants in *Mtb* or *Msm*.

## DISCUSSION

We have reported multiple lines of evidence that nargenicin acts as a DNA replication inhibitor in mycobacteria by targeting the essential DnaE1 polymerase, an enzyme identified recently as a highly vulnerable component of the DNA replication machinery in *Mtb* (Bosch *et al*., 2021). Unlike the commonly used nucleotide analogs that act as chain terminators through incorporation into the nascent DNA strand, nargenicin does not get incorporated into the DNA. Instead, it is wedged between the terminal base pair of the DNA substrate and the polymerase fingers domain, occupying both the position of the incoming nucleotide and the templating base, which is displaced by nargenicin. This binding mode is analogous to that of the human Pol α inhibitor, aphidicolin, which is derived from the fungus, *Cephalosporium aphidicola*, and unrelated in structure to nargenicin, indicating that these inhibitors have evolved independently. This unusual mechanism might explain the observation that the antimycobacterial activity of nargenicin was not diminished by over-expression of the cognate target, DnaE1. Based on this mechanism, the DnaE homologue in the *Nocardia* sp. CS682 producer organism would presumably need to bind nargenicin in a DNA-independent manner in order to fulfil its postulated “decoy” role in self-resistance (Pidot and Rizzacasa, 2020).

Nargenicin-mediated disruption of replisome function triggers a physiological response in *Mtb* which resembles that elicited by genotoxins which cause double-stranded breaks (DSBs) (mitomycin C, fluoroquinolones) (Boshoff *et al*., 2003). This features upregulation of genes encoding the recombinase involved in recombination repair (*recA*), the mutasome responsible for SOS mutagenesis and damage tolerance (*dnaE2*, *imuA’*, *imuB*) (Boshoff *et al*., 2003; Warner *et al*., 2010), the DSB-resecting motor-nuclease (*adnAB*) (Sinha et al., 2009) and a cell wall hydrolase (*chiZ*) (Chauhan et al., 2006), amongst other DNA-damage-responsive genes in mycobacteria. The DNA damage response to nargenicin begs the question of whether pharmacological inhibition of DnaE1 by this or other inhibitors might have the unintended consequence of inducing chromosomal mutations which could fuel the evolution of drug resistance, as documented for sub-lethal treatment of mycobacteria and other organisms by fluoroquinolones (Bush et al., 2020; Gillespie et al., 2005). This question is the subject of ongoing investigation in our laboratories. The concomitant downregulation of *ftsZ* (de Wet *et al*., 2020), *sepF* (Gupta et al., 2015), *whiB2* (Bush, 2018) and *ripA* (Gupta *et al*., 2015) is consistent with cellular elongation resulting from a block in cell division, followed by cell death. Ablation of the SOS response by deletion of *recA*, or a component thereof (mutasome function) by deletion of *dnaE2*, had no impact on the antimycobacterial activity of nargenicin, suggesting that the SOS-induced DNA repair, damage tolerance and mutagenesis systems are unable to rescue mycobacteria from the growth inhibitory effects of nargenicin. Instead, an arrest in cell division, as evidenced by bacillary elongation, appears to precede cell death. Another feature of the nargenicin mode of action was the late, strong signal elicited in the GFP release assay. The induction of *chiZ* and downregulation of *fbpC* might be telling in this regard: firstly, the damage-inducible protein, ChiZ (Burby and Simmons, 2020), been reported to arrest cell division, increase filamentation and induce cell lysis when overexpressed (Chauhan *et al*., 2006). Secondly, inactivation of the mycolyltransferase, FbpC, a member of the antigen 85 complex involved in the synthesis of trehalose dimycolate and mycolylarabinogalactan, which are key components of the mycobacterial cell envelope, has been shown to significantly reduce the mycolate content and increase the permeability of the cell envelope to small hydrophobic and hydrophilic molecules (Jackson et al., 1999). Thus, in addition to its replication-arresting activity, nargenicin may also compromise the integrity of the mycobacterial outer membrane and thus, act as a potentiator of other antitubercular agents whose efficacy is limited by permeation across the mycobacterial cell envelope. This intriguing possibility is the subject of active investigation in our laboratories.

In summary, the results reported here have positioned DnaE1 as a promising new TB drug target and laid the foundation for target-led drug discovery efforts focused on this enzyme.

## SIGNIFICANCE

The ongoing evolution and spread of drug-resistant strains of *Mycobacterium tuberculosis* underscores the importance of identifying and validating new tuberculosis (TB) drug targets. In this study, we report the chemical validation of one such target, the replicative DNA polymerase, DnaE1, with the narrow-spectrum antimicrobial agent, nargenicin. We show that nargenicin mediates its bactericidal activity against *M. tuberculosis* through interaction with DnaE1 in a manner that depends upon the presence of the DNA substrate. In this interaction, the nargenicin molecule wedges itself between DnaE1 and the terminal base pair of the DNA and occupies the place of both the incoming nucleotide and the templating base. By analysing the physiological consequences of *M. tuberculosis* exposure to nargenicin, we show that the arrest in bacillary replication resulting from the nargenicin-DnaE1 interaction triggers induction of a DNA damage response coupled with an arrest in cell division and an apparent weakening of the mycobacterial cell envelope. In addition to strongly reaffirming the value of natural products as a source of novel antitubercular agents, this work has provided the rationale and platform for focusing target-led drug discovery efforts on a promising new TB drug target.

## ACKNOWLEDGEMENTS

This work was supported by grants from the Bill & Melinda Gates Foundation (OPP1158806 to C.E.B.III and V.M.; INV-004757 to V.M. and INV-002474 to M.H.L.), the South African Medical Research Council, the National Research Foundation of South Africa, the Oppenheimer Memorial Trust, and an International Research Scholar’s grant from the HHMI (#55007649) (to V.M.), an African Career Accelerator Award from the Crick African Network (to M.K.M.), and, in part, by the Intramural Research Program of NIAID (to H.I.M.B. and C.E.B.III). We acknowledge use of the ilifu cloud computing facility (www.ilifu.ac.za) and thank Timothy de Wet for advice and assistance with morphotyping experimentation and analysis, Dirk Schnappinger and Jeremy Rock for kindly providing reagents, and Curtis Englehart for technical advice.

## AUTHOR CONTRIBUTIONS

M.D.C. and M.K.M. performed the microbiological profiling and RNA-seq experiments and analyses; G.L.A. and H.I.M.B. performed the DNA microarray and macromolecular incorporation assays; R.T. performed the polymerase assays; A.B. and M.H.L. performed the cryo-EM; J.A. provided bioinformatics support; and S.L. and Y.-M.A., provided technical support. The experiments were designed, and data analyzed, by M.D.C., M.K.M., G.L.A., B.M.C., D.B.O., D.F.W., C.E.B. III, H.I.M.B., M.H.L. and V.M. The manuscript was written by M.D.C., M.K.M., M.H.L. and V.M., and all authors read and edited it. M.H.L., D.B.O, C.E.B.III and V.M. were Team Leads.

## STAR METHODS

### RESOURCE AVAILABILITY

#### Lead Contact

Further information and requests for resources and reagents should be directed to and will be fulfilled by the lead contact, Valerie Mizrahi.

#### Materials availability

Plasmids and bacterial strains generated for this study are available upon request.

#### Data and code availability

RNA-seq datasets from this study are deposited in the NCBI Sequence Read Archive (SRA) repository (PRJNA722614) and are publicly available as of the date of publication. Atomic models and cryo-EM maps have been deposited to the Protein Data Bank and the Electron Microscopy Database under accession codes PDB XXX and EMD YYY. Accession numbers are listed in the **key resources table**. The paper does not report original code. The pipeline for RNA-seq analysis can be found at the GitHub repository (https://github.com/jambler24/bacterial_transcriptomics). Microscopy data are available from the lead contact upon request. Any additional information required to reanalyse the data reported in this paper is available from the lead contact upon request.

### EXPERIMENTAL MODEL AND SUBJECT DETAILS

#### Bacterial strains, culture conditions and media

The strains used in this study are listed in the **key resources table**. These include the parental wildtype strains, Mtb H37Rv (Ioerger et al., 2010) and *Msm* mc^2^155 (Snapper et al., 1990). Clinical isolates were obtained from samples collected from new TB cases and retreatment cases of subjects who were enrolled in a prospective longitudinal cohort study (ClinicalTrials.gov identifier, NCT00341601) at the National Masan Tuberculosis Hospital in the Republic of Korea from May 2005 to December 2006 (Shamputa et al., 2010). Mycobacterial strains were cultured in various media depending on the assay. 7H9 OADC was prepared by supplementing Middlebrook 7H9 (Difco) with 10% oleic acid-albumin-dextrose-catalase (OADC) enrichment (Difco), 0.2% glycerol and either 0.05% Tween-80 (7H9/OADC/Tw) or 0.05% Tyloxypol (7H9/OADC/Tx). 7H9/Glu/ADC/Tw medium was prepared by substituting 10% OADC with 10% albumin-dextrose-catalase (ADC) enrichment (Difco). Similarly, 7H9/Glu/CAS/Tx was prepared by supplementing 7H9 with 0.4% glucose, 0.03% casitone (CAS), 0.081% NaCl and 0.05% Tx. Glycerol-alanine-salts with iron (GAST-Fe/Tw) medium, pH 6.6, was prepared with 0.03% CAS, 0.005% ferric ammonium citrate, 0.4% dibasic potassium phosphate, 0.2% citric acid, 0.1% L-alanine, 0.12% MgCl_2_, 0.06% potassium sulphate, 0.2% ammonium chloride, 0.018% of a 1% sodium hydroxide solution, 1% glycerol and 0.05% Tween-80. GAST/Tw, an iron limiting media, was made as described above, but excluding ferric ammonium citrate. All *Mtb* cultures were incubated at 37°C in sealed culture flasks with no agitation. Cells were plated onto Middlebrook 7H10 agar plates with 7H10 agar base (Difco) supplemented with 10% OADC and 0.5% glycerol. Unless indicated otherwise, microbiological assays using the strains described below were performed in 7H9/OADC/Tw media.

The fluorescent reporter strain, H37Rv-GFP (Abrahams et al., 2012), and bioluminescent reporter strain P*recA*-LUX (Naran *et al*., 2016) were grown in media supplemented with kanamycin (Kan) at 20 µg/ml, whereas the *Mtb* mScarlet strain and *Msm* ΔL mutant were grown in media supplemented with hygromycin (Hyg) at 50 µg/ml. *Mtb* and *Msm* strains carrying the P*_UV15-Tet_*-*dnaE1*-*MYC*::L5 vector (Rock *et al*., 2015) were grown in media containing Kan at 50 µg/ml, and supplemented with ATc at 100ng/ml to induce expression of *dnaE1*. The inducible CRISPRi hypomorphs were grown in media containing Kan (25 µg/ml) and Hyg (50 µg/ml) and supplemented with ATc at 100ng/ml to induce transcriptional silencing. Minimal inhibitory concentrations (MICs) were determined against a range of clinical isolates: *Mtb* CDC1551 (Valway et al., 1998); *Mtb* HN878 (Manca et al., 2001); drug susceptible isolates, *Mtb* 0A029*, Mtb* 0A031 and *Mtb* 0B229; multi-drug resistant isolates, *Mtb* 0B123 (resistant to isoniazid (INH^R^), ofloxacin (OFX^R^), *para*-amino salicylic acid (PAS^R^), streptomycin (STR^R^), rifampicin (RIF^R^); *Mtb* 0A024 (ethambutol (EMB^R^), INH^R^, KAN^R^, PAS^R^, pyrazinamide (PZA^R^), STR^R^, ethionamide (ETH^R^), RIF^R^), *Mtb* 0B026 (EMB^R^, INH^R^, KAN^R^, PAS^R^, RIF^R^); and an extensively drug resistant strain, *Mtb* 0B014 (EMB^R^, INH^R^, KAN^R^, OFX^R^, PAS^R^, RIF^R^) (Shamputa *et al*., 2010).

### METHOD DETAILS

#### Drug susceptibility testing

MIC testing was performed by broth microdilution assay (Abrahams *et al*., 2012) and quantitatively analyzed with the colorimetric alamarBlue cell viability reagent (Thermo Fischer Scientific), as previously described (Chengalroyen *et al*., 2020).

#### Bioluminescence assay

P*recA*-LUX (Naran *et al*., 2016) was grown to an OD_600_ ~ 0.4, diluted 10-fold in 7H9/OADC/Tw and inoculated into white, clear-bottom, 96-well microtiter plates (Greiner CellStar^®^) containing a two-fold serial dilutions of drug. Plates were incubated at 37°C and luminescence recorded every 24 h for 8 days using a SpectraMax i3x plate reader (Molecular Devices). Data were plotted in Prism 9 (GraphPad).

#### GFP release assay

As described previously (Chengalroyen *et al*., 2020), H37Rv-GFP was grown to an OD_600_ of ~ 0.3 in 7H9 OADC and exposed to drug at 1× or 10× MIC. Every 24 h, over a period of 8 days, 200 µl of culture was harvested, pelleted by centrifugation, and the supernatant transferred to a black, clear-bottom 96-well microtitre plate (Greiner CellStar^®^) and fluorescence (excitation, 540 nm; emission, 590 nm) measured using a SpectraMax i3x plate reader (Molecular Devices). Fluorescence intensity was normalized by OD_650_ and standardized to the value of the drug-free control for each sample.

#### Time-kill kinetics

*Mtb* was inoculated in culture medium at an OD_600_ of 0.002 and drug added at a concentration of either 1×, 5× or 10× MIC. Cultures were incubated in sealed culture flasks and 1 ml aliquots harvested every 24 h over 8 days. The samples were washed twice in fresh media. One hundred µl aliquots of 10-fold serial dilutions were plated of 7H11 agar and colony forming units (CFUs) enumerated after incubation for 3-4 weeks.

#### Macromolecular incorporation assays

Macromolecular incorporation assays were performed as described (Barrow et al., 2003; Cotsonas King and Wu, 2009). Briefly, *Mtb* cultures were grown to early exponential phase (OD_600_ ~ 0.3) and 1 μCi/ml [^3^H]-uracil, 2.5 μCi/ml [3H]-phenylalanine, 10 μCi/ml [^3^H]-N-acetyl glucosamine, and 1 μCi/ml [^14^C]-acetate added to quantify the incorporation of the radiolabeled precursors into either total nucleic acid (i.e., DNA and RNA), protein, cell wall, and fatty acids, respectively. Cells were incubated at 37°C for 1 h and 150 μl transferred to 96-well microtiter plates containing 150 μl of each test compound. Nargenicin was used at 2× and 20× MIC with 1% DMSO included as an untreated control. The specificity of assays was monitored by the inclusion of the pathway-specific antibiotics OFX (5 μg/mL), STR (10 μg/mL), D-cycloserine (DCS, 5 μg/mL), and INH (0.2 μg/mL) as positive controls. The assay plates were incubated at 37°C for 24 h and precursor incorporation terminated by the addition of 300 μl of 20% trichloroacetic acid (TCA). The samples were incubated at 4°C for 1 h and the precipitates collected by vacuum filtration with a 96-well MultiScreen GFC glass fiber plate (Millipore). Precipitates were washed three times with 10 % TCA followed by three 95% ethanol washes and the plates allowed to air dry. Precipitates were resuspended in 50 μl MicroScint 20 (PerkinElmer) and the radioactivity on each filter measured in a MicroBeta Liquid Scintillation Counter (PerkinElmer). To distinguish between the incorporation of [^3^H]-uracil into DNA vs. RNA, the RNA was hydrolyzed with 500 μl of 1M KOH at 37°C for 16 h and neutralized with 125 μl HCl. Samples were then precipitated by adding 625 μl 20% TCA and the amount of residual radioactivity present in the DNA precipitates quantified following filtration and washing as described above. All samples were analyzed in duplicate, and results represent the percentage of radiolabel incorporation relative to the DMSO-treated control from two independent replicates.

#### Microscopy

*Msm* bacilli were imaged to determine their terminal phenotypes under exposure to varying concentrations of antibiotic as previously described (de Wet *et al*., 2020). Strains were grown to late-log phase (OD_600_ ~ 0.8), filtered once through a Millex syringe filter (5 µm pore size, Millipore) and diluted 1:40 into fresh media. Samples were left untreated, exposed to carrier (DMSO only), or to varying concentrations of nargenicin in DMSO (1× MIC, 2× MIC, 4× MIC) and incubated for 18 h at 37°C while shaking. After exposure cultures were spotted onto low-melt agarose pads and imaged on a ZEISS Axio Observer using a 100×, 1.4 na Objective with Phase Contrast and Colibri 7 fluorescent illumination system. Images were captured using a Zeiss Axiocam 503. Image processing, cell measurements and analysis were performed in the FIJI Plugin MicrobeJ (Ducret et al., 2016; Schindelin et al., 2012), R (R Core Team, 2020; RStudio Team, 2020) and UMAP as described (de Wet *et al*., 2020).

#### Transcriptional profiling

Microarray experiments and analyses were performed by the NIAID Microarray Research Facility, as previously described (Boshoff *et al*., 2003), including two independent samples for each treatment condition. Datasets from cultures exposed to mitomycin C (0.2 µg/ml), and levofloxacin (10 µg/ml) were compared to nargenicin (129 µg/ml). The top 300 upregulated or downregulated genes, ranked by the average Log2 fold-change in expression data from two biological repeats, were compared to generate gene shortlists common to all three treatments.

For RNA-seq, qRT-PCR and microarray experiments, *Mtb* cultures (20-30 ml) were grown either in roller bottles or in culture flasks on a shaker to mid-exponential phase (OD_600_ ~ 0.3-0.5) prior to treatment with nargenicin at 1× or 10× MIC for 6 h. Cells were harvested by centrifugation at 3000 × *g* for 10 min and resuspended in 1 ml Qiazol Lysis Reagent (Qiagen). Cells were lysed with 0.1 mm Zirconia/Silica beads (BioSpec) in a MagNA Lyser Homogenizer (Roche) (6000 rpm, 30 s) three times with 1 min cooling intervals. Samples were centrifuged at 10 000 × g for 5 min at 4°C and the supernatant transferred into a clean tube containing an equal volume of 100% ethanol. The RNA was purified and treated with DNase on-column using the Direct-zol RNA MiniPrep kit (Zymo Research) according to the manufacturer’s protocol. Samples were eluted in 50 μl of RNase- and DNase-free water. Purified RNA was treated with DNase for an additional 60 min at 37°C using the TURBO DNA-free kit (Ambion) according to the manufacturer’s protocol. In preparation for microarray analysis and RNA-seq, the sample quality was confirmed using a Bioanalyzer RNA 6000 Nano Kit and Chips (Agilent). For RNA-seq experiments, three independent biological replicates of both nargenicin-treated (10× MIC) and untreated samples were performed. Library preparation and sequencing was done by Admera Health (NJ, USA) using the Illumina NovaSeq S4 sequencing platform. The sequencing strategy included an average of 60 million 150 bp paired end reads per sample. Reads were demultiplexed to generate raw fastq files for each sample and data deposited in the NCBI SRA repository (PRJNA722614). Initial quality control (QC) of the raw fastQ files was performed using FastQC (Andrews, 2010). Reads were trimmed and adapters removed using Trim Galore. Further QC was done by aligning reads using BWA to the reference genome of *Mtb* H37Rv, ASM19595v2, GenBank assembly accession no. GCA_000195955.2 (https://www.ncbi.nlm.nih.gov/assembly/GCF_000195955.2), running RSeQC (Wang et al., 2016) and dupRadar (Sayols et al., 2016), and an amalgamated report generated using MultiQC (Ewels et al., 2016). Transcript quantification was performed using Salmon in mapping-based mode (Patro et al., 2017). Normalization and differential expression analysis were done using DESeq2 (Love et al., 2014) with count normalization by DESeq2’s median or ratios. *P*-values were adjusted for multiple-testing using the Benjamini-Hochberg approach, and genes which displayed an absolute Log2 fold-change > 1 and an adjusted *p*-value < 0.05 were considered differentially expressed. Data were visualized in R and functional enrichment of upregulated and downregulated shortlists as compared to the full genome was performed in STRING (Szklarczyk et al., 2018) using Gene Ontologies, STRING local network clusters, annotated keywords, KEGG pathways and InterPro protein domains and features as categories. Multiple comparisons were compensated for using the false discovery rate (FDR), with significant enrichment considered as FDR > 0.05.

For qRT-PCR experiments, following TURBO DNase treatment, 250 ng of the RNA was converted to cDNA using SuperScript® IV Reverse Transcriptase (Thermo Fischer Scientific). Regions of interest were amplified using primer pairs described in **Table S3** and Power SYBR® Green PCR master mix (Thermo Fischer Scientific) and transcript levels for three independent samples quantified on a PikoReal real-time PCR system (Thermo Fischer Scientific). Transcript levels of target genes were normalized to *sigA*.

#### Construction of fluorescent *dnaE1* hypomorphs

The ATc-regulated CRISPRi system developed by Rock *et al*. (2017) was used to construct inducible *dnaE1*-targeting *Mtb* hypomorphs carrying the mScarlet fluorescence reporter (Kolbe et al., 2020) (see **key resources table**). Briefly, two oligonucleotides complementary to the *dnaE1* targeting sequence (**Table S3**) were annealed and cloned in pLJR965, and the presence of the sgRNA confirmed by Sanger sequencing. The sequence-verified constructs were electroporated into *Mtb* mScarlet, selecting on media supplemented with Kan (25 µg/ml) and Hyg (50 µg/ml).

#### Drug susceptibility testing using hypomorphs

To assess the impact of *dnaE1* silencing on drug susceptibility, the hypomorphs and vector control strains were grown to an OD_600_ of 1.0 and diluted to an OD_600_ of 0.01 in media either with ATc (200 ng/ml) or without the inducer. Fifty µl of the diluted culture was inoculated into each well of a MIC plate containing 50 µl of media with 2-fold dilutions of drug. Microtitre plates were incubated at 37°C for 14 days and the fluorescence (594 nm, excitation; and 569 nm, emission) recorded using a Spectramax i3x plate reader. Each strain was normalized to the no-drug control to determine the percentage growth inhibition as a function of drug concentration. Dose-response curves were plotted in Prism 9 (GraphPad).

#### Protein expression and purification

*Mtb* DnaE1 was expressed in *Msm* and purified as previously described (Rock *et al*., 2015). *S. aureus* DnaE and *E. coli* Pol IIIα were expressed in *E. coli* BL21 and purified as previously described (Painter *et al*., 2015; Toste Rego et al., 2013).

#### DNA polymerase assay

DNA polymerase activity was measured using a real-time polymerase assay as described previously (Rock *et al*., 2015). Briefly, reactions were performed using 5 nM DNA polymerase, 10 nM of fluorescently labelled DNA substrate (Primer: 5’-TAGGACGAAGGACTCCCAACTTTAGGTGCG, Template: 6-FAM-5’-CCCCCCCCCATGCATGCGCACCTAAAGTTGGGAGTCCTTCGTCCTA) and 100 nM of unlabeled DNA substrate (same sequence as above). Reactions contained 100 μM of each dNTP, 5 mM MgSO_4_, 50 mM HEPES pH 7.5, 100 mM potassium glutamate, 2 mM DTT, 0.5 mg/ml BSA and 10 nM - 10 μM nargenicin. 10 μL reactions were measured for 20 minutes at 24 °C in a 384-well plate using a Clariostar plate reader (BMG LABTECH) with excitation and emission filters at 485 and 520 nm, respectively.

#### Fluorescence anisotropy

DNA binding was measured using 5 nM of a Cy3-labelled DNA substrate (Primer: Cy3-5’-GGTAACGCCAGGGTTTTCCCAGTC3, Template 5’-CGCTCACTGGCCGTCGTTTTACAACGTCGTGACTGGGAAAACCCTGGCGTTACC) and 1 nM - 40 μM DNA polymerase. Reactions conditions contained 25 mM HEPES pH 7.5, 50 mM potassium glutamate, 2 mM DTT, 0.5 mg/ml BSA. 10 μL reactions were measured at 24 °C in a 384-well plate using a Clariostar plate reader with excitation and emission filters at 540 and 590 nm, respectively.

#### Cryo-EM sample preparation and imaging

Purified *Mtb* DnaE1 was diluted to 4 μM in 20 mM PIPES pH 7.0, 50 mM potassium glutamate, 5 mM MgCl2, 2 mM DTT, and 0.01% Tween-20. The diluted protein was incubated for 5 min with 20 μM DNA substrate (Template: 5’-GATAGAGCAGAAGGACGAAGGACTCCCAACTTTAGGTG, Primer: 5’-GCACCTAAAGTTGGGAGTCCTTCGTCCT*T, where the asterisk marks the position of a phosphorothioate bond). Three μl of sample were adsorbed onto glow-discharged copper R2/1 holey carbon grids (Quantifoil). Grids were glow discharged 45 seconds at 25 mA using an EMITECH K950 apparatus. Grids were blotted for one second at ~80% humidity at 4°C and flash frozen in liquid ethane using a Leica EM GP plunge freezer. The grids were loaded into a Titan Krios (FEI) electron microscope operating at 300 kV with a Gatan K3 detector. The slit width of the energy filter was set to 20 eV. Images were recorded with EPU software (Thermo Fisher Scientific) in counting mode. Dose, magnification, and pixel size are detailed in **Table 1**.

**Table 1.**
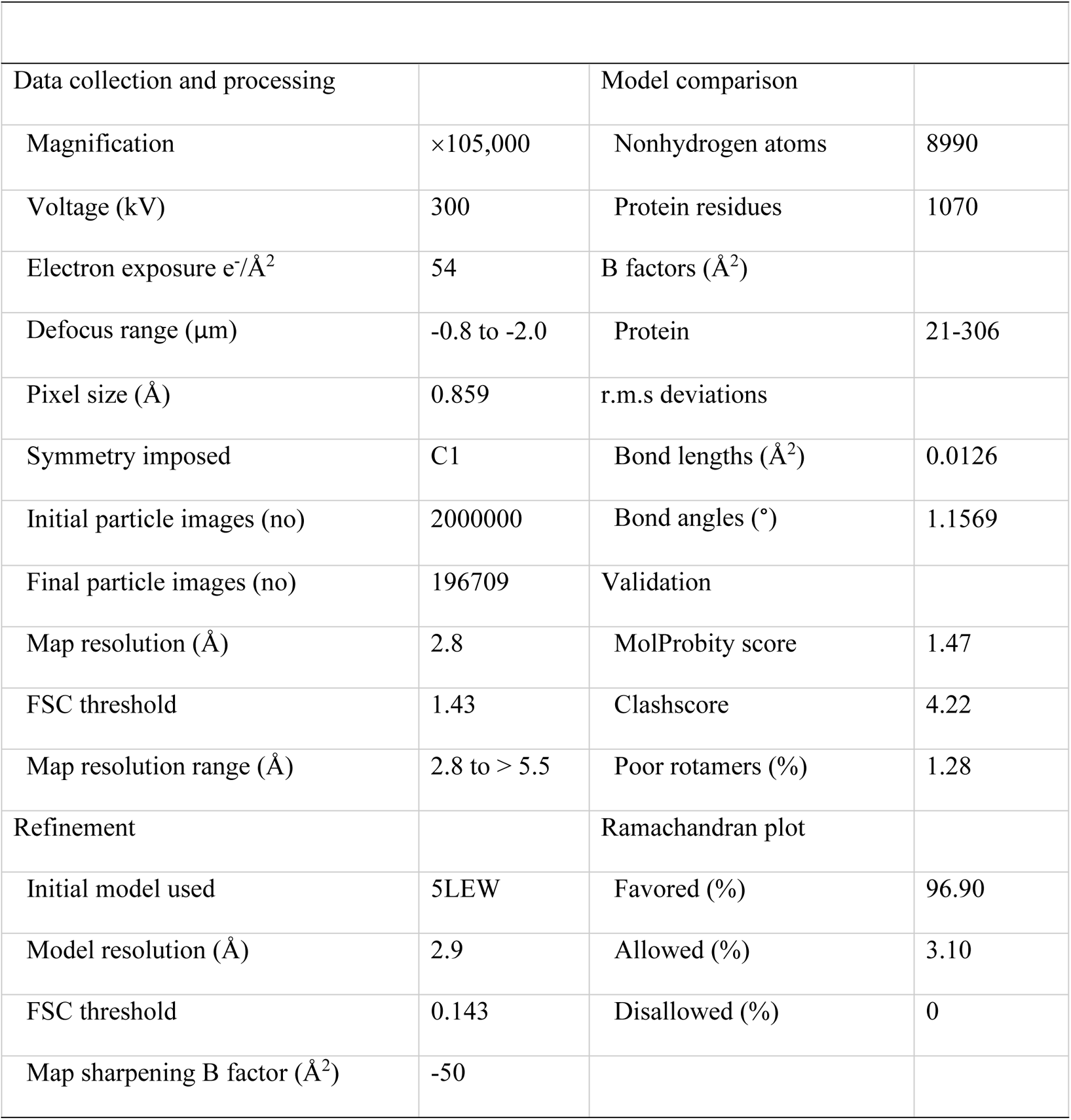
Cryo-EM data collection, refinement, and validation statistics.

| **Table 1. Cryo-EM data collection, refinement, and validation statistics** |  |  |  |
| --- | --- | --- | --- |
| Data collection and processing |  | Model comparison |  |
| Magnification | ×105,000 | Nonhydrogen atoms | 8990 |
| Voltage (kV) | 300 | Protein residues | 1070 |
| Electron exposure $e^-/\text{\AA}^2$ | 54 | B factors ($\text{\AA}^2$) | |
| Defocus range ($\mu\text{m}$) | -0.8 to -2.0 | Protein | 21-306 |
| Pixel size ($\text{\AA}$) | 0.859 | r.m.s deviations | |
| Symmetry imposed | C1 | Bond lengths ($\text{\AA}^2$) | 0.0126 |
| Initial particle images (no) | 2000000 | Bond angles ($^\circ$) | 1.1569 |
| Final particle images (no) | 196709 | Validation |  |
| Map resolution ($\text{\AA}$) | 2.8 | MolProbity score | 1.47 |
| FSC threshold | 1.43 | Clashscore | 4.22 |
| Map resolution range ($\text{\AA}$) | 2.8 to > 5.5 | Poor rotamers (%) | 1.28 |
| Refinement |  | Ramachandran plot |  |
| Initial model used | 5LEW | Favored (%) | 96.90 |
| Model resolution ($\text{\AA}$) | 2.9 | Allowed (%) | 3.10 |
| FSC threshold | 0.143 | Disallowed (%) | 0 |
| Map sharpening B factor ($\text{\AA}^2$) | -50 | | |

#### Cryo-EM image processing

All image processing was performed using RELION 3.1 (Zivanov et al., 2018). The images were drift corrected using RELION’s own (CPU-based) implementation of the UCSF motioncor2, and defocus was estimated using gCTF (Zhang, 2016). LoG-based auto-picking was performed on all micrographs and picked particles were 2D classified. After three rounds of 2D classification, classes with different orientations were selected for initial model generation in RELION. The initial model was used as reference for 3D classification into different classes. The selected classes from 3D classification were subjected to 3D auto refinement followed by different rounds of CTF refinement plus a final round of Bayesian polishing. Polished particles were used for 3D auto-refine job and the final map was post-processed to correct for modulation transfer function of the detector and sharpened by applying a negative B-factor manually set to −50. A soft mask was applied during post processing to generate FSC curves to yield a map of average resolution of 2.9 Å. The RELION post-processed map was used to generate improved-resolution EM maps using the SuperEM method (Subramaniya et al., 2021), which aided in model building and refinement. Model building was performed using Coot (Emsley et al., 2010), REFMAC5 (Murshudov et al., 2011), the CCPEM-suite (Nicholls et al., 2018) and Phenix (Liebschner et al., 2019). Details on model refinement and validation are shown in **Table 1**. In brief, model building started by rigid-body fitting of the known DnaE1 crystal structure (PDB 5LEW) (Baños-Mateos *et al*., 2017) into experimental density map using Coot. The DNA molecule was generated, and rigid body fitted into experimental density map using Coot. Next, we carried out one round of refinement in REFMAC5 using jelly-body restraints, and the model was further manually adjusted in Coot. Final refinement and model validation were performed using Phenix.

#### Quantification and statistical analysis

Statistical details are given in methods sections and figure legends, these include details of the experiments, numbers of replicates (technical and/or experimental), statistical software used and thresholds of significance. Significance was generally determined as p<0.05 and correction for multiple comparisons was performed, as appropriate. Independent experiments were performed a minimum of two times and these data were utilised for the generation of summary statistics (mean and standard deviation). Replicate data are included within each figure, as indicated in figure legends, else data are described as a representative experiment. In addition, DNA polymerase assays and DNA binding experiments were performed in three or more independent experiments. Data were not excluded from experimental datasets prior to or during analyses other than during cryo-EM data processing, where particles that did not possess high resolution features were removed following standard procedures for cryo-EM structure determination.

### SUPPLEMENTAL ITEMS

Document S1. Figures S1-S6 and Tables S1-S3

Data S1. Dataset S1 comprising output genelists of RNA-seq differential gene expression analysis and STRING functional analysis of significantly differentially expressed genes. Related to Figure 2 and Figure 2C.

## Key resources table

| REAGENT or RESOURCE | SOURCE | IDENTIFIER |
| --- | --- | --- |
| Antibodies |  |  |
| Bacterial and virus strains |  |  |
| *Escherichia coli* DH5α | Thermo Fisher Scientific | Cat#18265017 |
| *Escherichia coli* BL21 | New England Biolabs | Cat#C2530H |
| *Mycobacterium smegmatis* mc<sup>2</sup>155 | (Snapper *et al.*, 1990) | N/A |
| *Mycobacterium tuberculosis* H37Rv (H37RvMA) | (Ioerger *et al.*, 2010) | N/A |
| *Mycobacterium tuberculosis* CDC1551 | Clinical isolate (Valway *et al.*, 1998) | N/A |
| *Mycobacterium tuberculosis* HN878 | Clinical isolate (Manca *et al.*, 2001) | N/A |
| *Mycobacterium tuberculosis* K04b00DS | Clinical isolate (Shamputa *et al.*, 2010) | N/A |
| *Mycobacterium tuberculosis* NIH_KA31b00DS | Clinical isolate (Shamputa *et al.*, 2010) | N/A |
| *Mycobacterium tuberculosis* K14b00DS | Clinical isolate (Shamputa *et al.*, 2010) | N/A |
| *Mycobacterium tuberculosis* K29b00MR | Clinical isolate (Shamputa *et al.*, 2010) | N/A |
| *Mycobacterium tuberculosis* K33b00MR | Clinical isolate (Shamputa *et al.*, 2010) | N/A |
| *Mycobacterium tuberculosis* K37b00XR | Clinical isolate (Shamputa *et al.*, 2010) | N/A |
| *Mycobacterium tuberculosis* K32b00MR | Clinical isolate (Shamputa *et al.*, 2010) | N/A |
| Biological samples |  |  |
| Chemicals, peptides, and recombinant proteins |  |  |
| Nargenicin A1, Antibiotic agent | Abcam | Cat#AB144312 |
| Griselimycin, Antibiotic agent | A gift from Prof. Rolf Müller & Dr. Jennifer Herrmann, Helmholtz Institute for Pharmaceutical Research Saarland | N/A |
| Levofloxacin, Antibiotic agent | Sigma-Aldrich (Merck) | Cat#28266 |
| Ofloxacin, Antibiotic agent | Sigma-Aldrich (Merck) | Cat#O8757 |
| Ciprofloxacin, Antibiotic agent | Sigma-Aldrich (Merck) | Cat#17850 |
| Isoniazid, Antibiotic agent |  |  |
| Mitomycin C | Sigma-Aldrich (Merck) | Cat# 10107409001 |
| Streptomycin, Antibiotic agent | Sigma-Aldrich (Merck) | Cat#85886 |
| D-cycloserine, Antibiotic agent | Sigma-Aldrich (Merck) | Cat#C6880 |
| Aphidicolin, Antibiotic agent | Sigma-Aldrich (Merck) | Cat#89458 |
| [6-<sup>3</sup>H]-uracil | Moravek | Cat#MT656 |
| L-[2,3,4,5,6-<sup>3</sup>H]-phenylalanine | American Radiolabeled Chemicals | Cat#ART1546 |
| [1,6-<sup>3</sup>H]-N-acetyl-D-glucosamine | American Radiolabeled Chemicals | Cat#ART0142 |
| [1-<sup>14</sup>C]-acetic acid, sodium salt | Moravek | Cat#MC125 |
| Critical commercial assays |  |  |
| alamarBlue™ Cell Viability Reagent | Thermo Fisher Scientific | Cat#DAL1100 |
| Qiazol Lysis Reagent | Qiagen | Cat#79306 |
| Direct-zol RNA MiniPrep kit | Zymo Research | Cat#R2050 |
| TURBO DNA-free kit | Ambion | Cat#AM1907 |
| Bioanalyzer RNA 6000 Nano Kit and Chips | Agilent | Cat#5067-1511 |
| SuperScript® IV Reverse Transcriptase | Thermo Fischer Scientific | Cat#18090010 |
| Power SYBR® Green PCR master mix | Thermo Fischer Scientific | Cat#4367659 |
| Deposited data |  |  |
| Transcriptomics datasets | This paper | NCBI SRA: PRJNA722614 |
| Atomic models | This paper | Protein Data Bank: PDB XXX (pending) |
| Cryo-EM maps | This paper | Electron Microscopy Database: EMD YYY (pending) |
| Experimental models: Cell lines |  |  |
| Experimental models: Organisms/strains |  |  |
| *Mycobacterium smegmatis* mc<sup>2</sup>155 Δ*recA* | (Machowski *et al.*, 2007) | N/A |
| *Mycobacterium smegmatis* mc<sup>2</sup>155 Δ*dnaE2* | (Warner *et al.*, 2010) | N/A |
| *Mycobacterium smegmatis* mc<sup>2</sup>155 P<sub>UV15-Tet</sub>-*dnaE1*-MYC::L5 | (Rock *et al.*, 2015) | N/A |
| *Mycobacterium tuberculosis* H37Rv-GFP | (Abrahams *et al.*, 2012) | N/A |
| *Mycobacterium tuberculosis* H37Rv PrecA-LUX | (Naran *et al.*, 2016) | N/A |
| *Mycobacterium tuberculosis* H37Rv P<sub>UV15-Tet</sub>-*dnaE1*-MYC::L5 | This paper | N/A |
| *Mycobacterium tuberculosis* H37Rv Δ*dnaE2* | (Boshoff *et al.*, 2003) | N/A |
| *Mycobacterium tuberculosis* H37Rv mScarlet | This paper | N/A |
| *Mycobacterium tuberculosis* H37Rv *dnaE1* sgRNA 3 | This paper | N/A |
| *Mycobacterium tuberculosis* H37Rv *dnaE1* sgRNA 6 | This paper | N/A |
| *Mycobacterium tuberculosis* H37Rv *dnaE1* sgRNA 11 | This paper | N/A |
| *Mycobacterium tuberculosis* H37Rv *dnaE1* sgRNA 13 | This paper | N/A |
| Oligonucleotides |  |  |
| See Table S3 for information on primers and other oligonucleotides used in this study |  |  |
| Recombinant DNA |  |  |
| P<sub>UV15-Tet</sub>-*dnaE1*-MYC::L5 | (Rock *et al.*, 2015) | N/A |
| pLJR965 | (Rock *et al.*, 2017) | N/A |
| Giles attB: P<sub>left</sub>* mScarlet | (Kolbe *et al.*, 2020) | N/A |
| pUC19-GI | (Kolbe *et al.*, 2020) | N/A |
| pLJR965_*dnaE1*_sgRNA3 | This paper | N/A |
| pLJR965_*dnaE1*_sgRNA6 | This paper | N/A |
| pLJR965_*dnaE1*_sgRNA11 | This paper | N/A |
| pLJR965_*dnaE1*_sgRNA13 | This paper | N/A |
| Software and algorithms |  |  |
| Microsoft Excel | Microsoft | N/A |
| Prism 9 | Graph Pad | N/A |
| SoftMax Pro Software | Molecular Devices | N/A |
| Clariostar Microplate Reader Software | BMG LABTECH | N/A |
| ZEN (blue) Imaging Software | ZEISS | N/A |
| FIJI | (Schindelin *et al.*, 2012) | https://fiji.sc |
| Microbe J Plugin | (Ducret *et al.*, 2016) | https://www.microbej.com |
| Spatial UMAP Analysis | (de Wet *et al.*, 2020) | https://osf.io/pdcw2/ |
| R ver. 4.1.0 | (R Core Team, 2020) | https://www.r-project.org/ |
| RStudio ver. 1.4.1717 | (RStudio Team, 2020) | https://www.rstudio.com |
| FastQC | (Andrews, 2010) | https://www.bioinformatics.babraham.ac.uk/projects/fastqc/ |
| TrimGalore | N/A | https://www.bioinformatics.babraham.ac.uk/projects/trim_galore/ |
| RSeQC | (Wang *et al.*, 2016) | http://rseqc.sourceforge.net |
| dupRadar | (Sayols *et al.*, 2016) | https://bioconductor.org/packages/release/bioc/html/dupRadar.html |
| MultiQC | (Ewels *et al.*, 2016) | https://multiqc.info |
| Salmon | (Patro *et al.*, 2017) | https://combine-lab.github.io/salmon/ |
| DESeq2 | (Love *et al.*, 2014) | https://bioconductor.org/packages/release/bioc/html/DESeq2.html |
| STRING ver. 11.0b | (Szklarczyk *et al.*, 2018) | https://string-db.org/ |
| EPU (Cryo-EM Single Particle Analysis Software) | Thermo Fisher Scientific | N/A |
| RELION 3.1 | (Zivanov *et al.*, 2018) | N/A |
| gCTF v1.06 | (Zhang, 2016) | N/A |
| Coot | (Emsley *et al.*, 2010) | https://pemsley.github.io/coot/blog/2021/02/02/release-0.9.4.1.html |
| REFMAC5 | (Murshudov *et al.*, 2011) | https://www2.mrc-lmb.cam.ac.uk/groups/murshudov/content/refmac/refmac.html |
| CCPEM-suite | (Nicholls *et al.*, 2018) |  |
| Phenix | (Liebschner *et al.*, 2019). | http://www.phenix-online.org |
| PyMOL | Schrödinger | https://pymol.org/ |
| Other |  |  |

