## Supplemental Information for "DNA-dependent binding of nargenicin to DnaE1 inhibits replication in *Mycobacterium tuberculosis*"

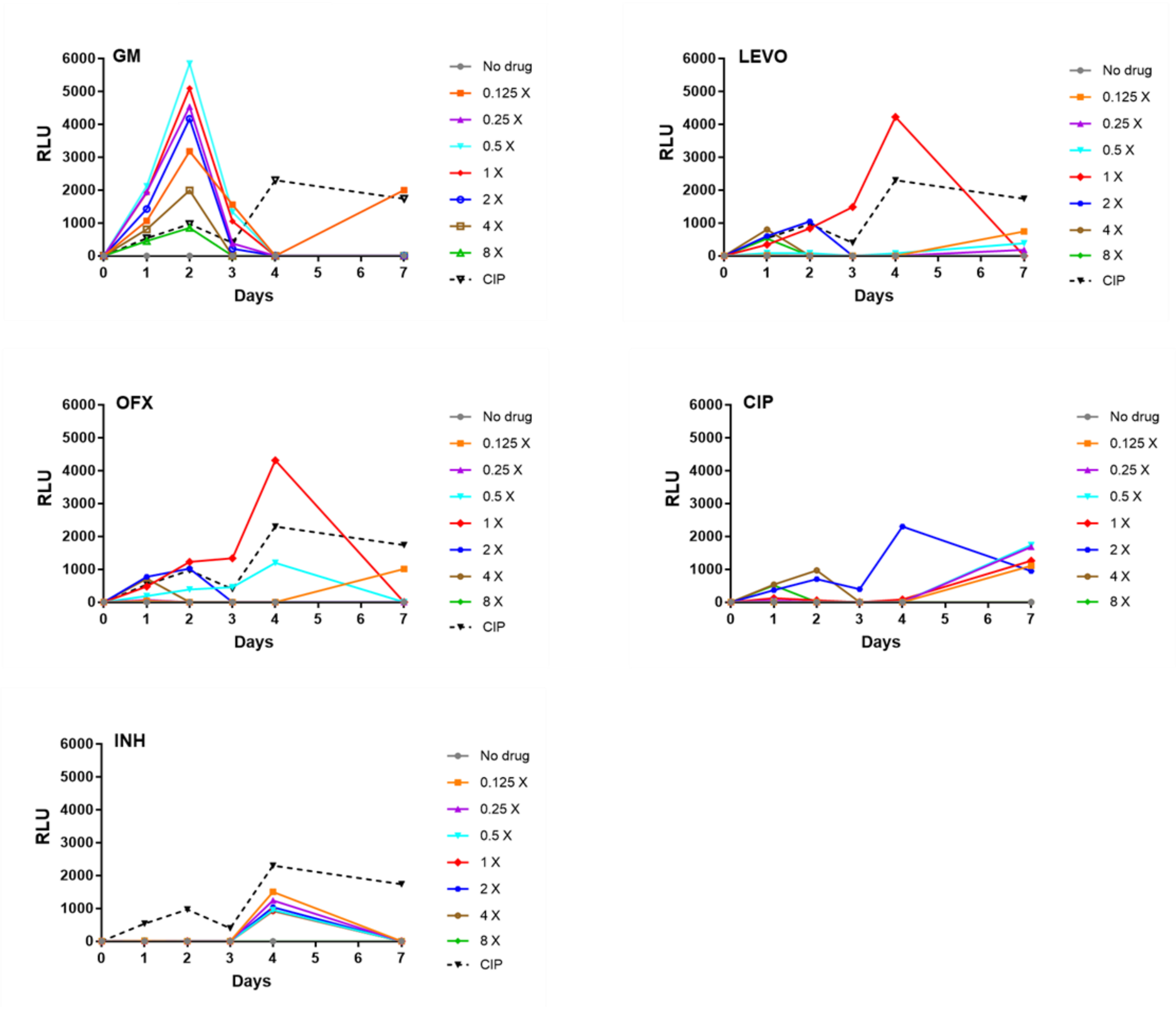


**Figure S1. Induction of *recA* bioluminescent reporter, PrecA-LUX (Naran et al., 2016), in response to various antitubercular agents. Related to Figure 2A.** Griselimycin (GM), levofloxacin (LEVO) and ofloxacin (OFX) were tested at concentrations ranging from 0.125× MIC to 8× MIC, and bioluminescence monitored over 7 days. Ciprofloxacin (CIP) was used as the positive drug control at 3.13 µM (2× MIC) and INH as the negative control. Data represent the average of two technical replicates from one representative experiment. Experiments were performed in duplicate.

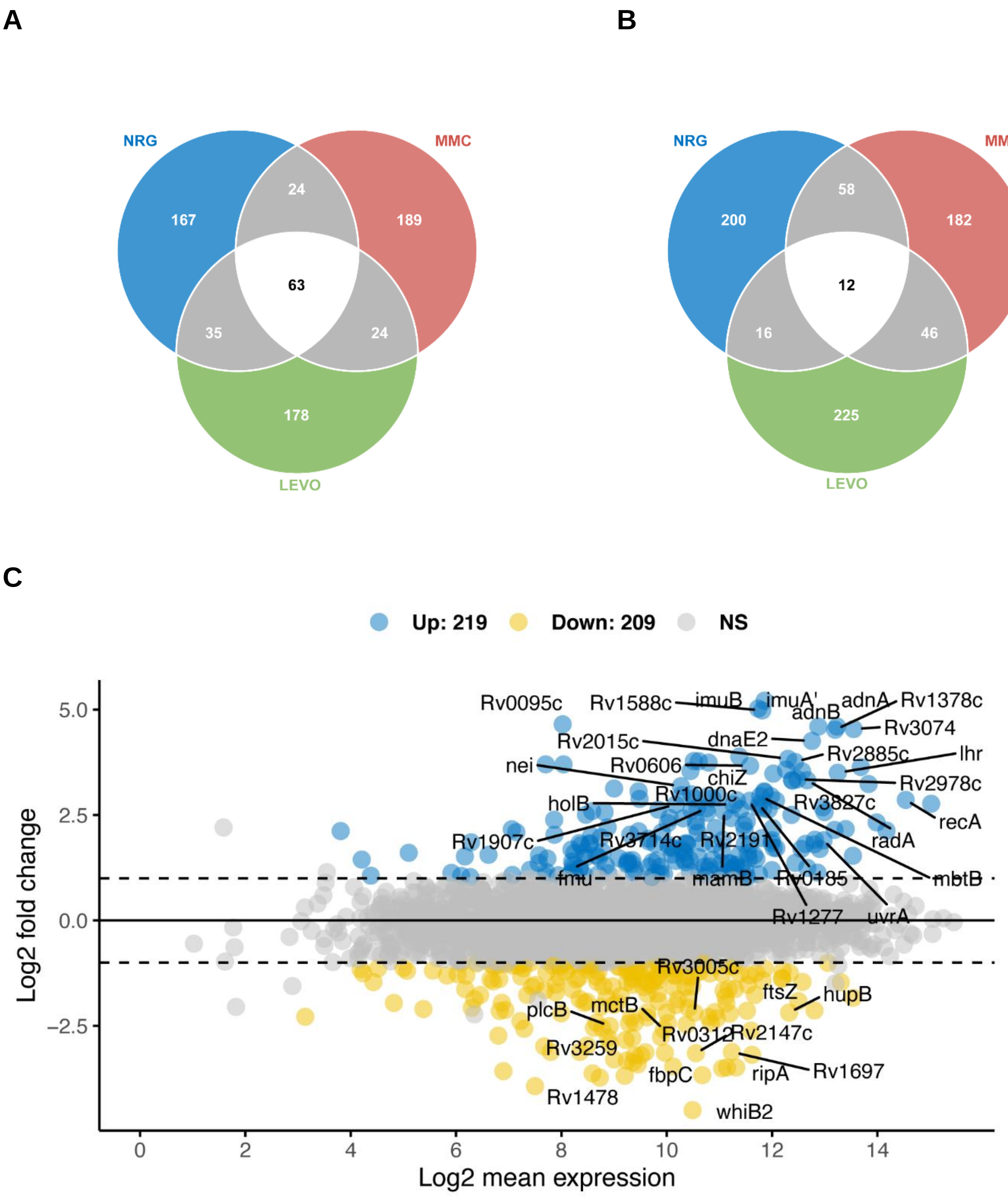


**Figure S2. Comparison of the transcriptional response of *Mtb* to nargenicin *versus* other genotoxins. Related to Figure 2B, and Data S1.** Venn diagram of the transcriptional response of *Mtb* determined by DNA microarray analysis illustrating the overlap of genes that are either (A) upregulated or (B) downregulated in response to nargenicin (NRG), mitomycin C (MMC) or levofloxacin (LEVO). The genes compared between the three datasets represent the top 300 genes ranked by the average Log2 fold change (FC) in expression data of two biological repeats. (C) Log ratios of RNA-seq data plotted against mean average expression (MA plot) of each gene to visualise the transcriptional response to nargenicin (20× MIC) with genes that show significant upregulation in blue and those that are significantly down regulated in yellow (FDR < 0.05, FC > 2). The genes are plotted in Figure S3 and listed in the Supplementary Data.

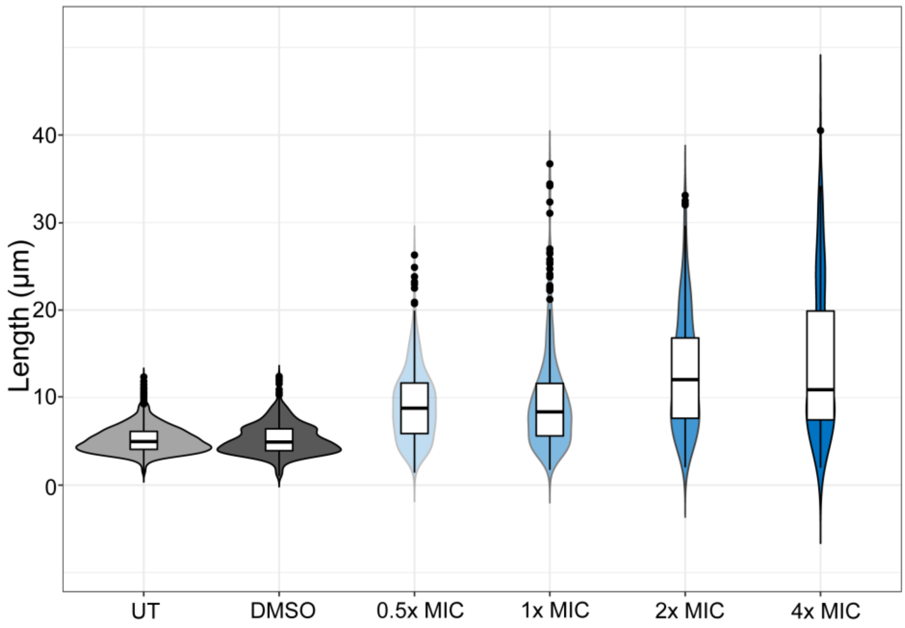


**Figure S3. Violin plots and boxplots of single-cell length data of from a population of *Msm* bacilli in response to treatment with varying concentrations of nargenicin. Related to Figure 2C.** UT, untreated control; DMSO, carrier-only control. Data for each sample were pooled from two independent repeats.

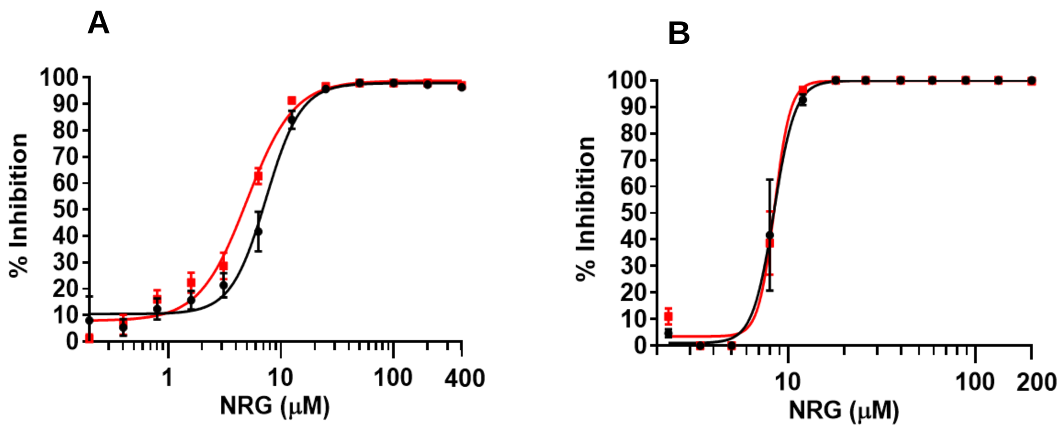


C

| Strain | *Msm dnaE1* | *Mtb dnaE1* |
|---|---|---|
| mc$^2$155 | 1.0 | - |
| mc$^2$155 P$_{UV15\text{-}Tet}$-*dnaE1-MYC*::*L5* -ATc | 1.2 | - |
| mc$^2$155 P$_{UV15\text{-}Tet}$-*dnaE1-MYC*::*L5* +ATc | 8.5 | - |
| H37Rv | - | 1.0 |
| H37Rv P$_{UV15\text{-}Tet}$-*dnaE1-MYC*::*L5* -ATc | 0.08 | 1.34 |
| H37Rv P$_{UV15\text{-}Tet}$-*dnaE1-MYC*::*L5* +ATc | 2.1 | 1.3 |

**Figure S4. Effect of *Msm dnaE1* overexpression on susceptibility of mycobacteria to nargenicin. Related to Figure 3C.** The recombinant strains (A) mc$^2$155 P$_{UV15\text{-}Tet}$-*dnaE1-MYC*::*L5* and (B) H37Rv P$_{UV15\text{-}Tet}$-*dnaE1-MYC*::*L5* were grown either in the absence (black) or presence (red) of 100 ng/ml ATc and assessed for NRG susceptibility. Bacterial growth inhibition was determined using the fluorescence-based resazurin assay. Means are calculated from three independent biological replicates. (C) Conditional over-expression of *Msm dnaE1* in *Msm* or *Mtb*, assessed by qRT-PCR analysis. Within each strain, quantification of the target *dnaE1* transcript was normalized to the level of endogenous *sigA* transcript. Normalized *dnaE1* transcript levels were divided by the level of expression of the endogenous *dnaE1* transcript to determine relative transcript levels. Comparison of relative transcript levels in the -ATc vs. +ATc samples of the recombinant strains confirmed ATc-dependent induction of *Msm dnaE1* in both *Msm* and *Mtb*. The data represent the results from three independent repeats.

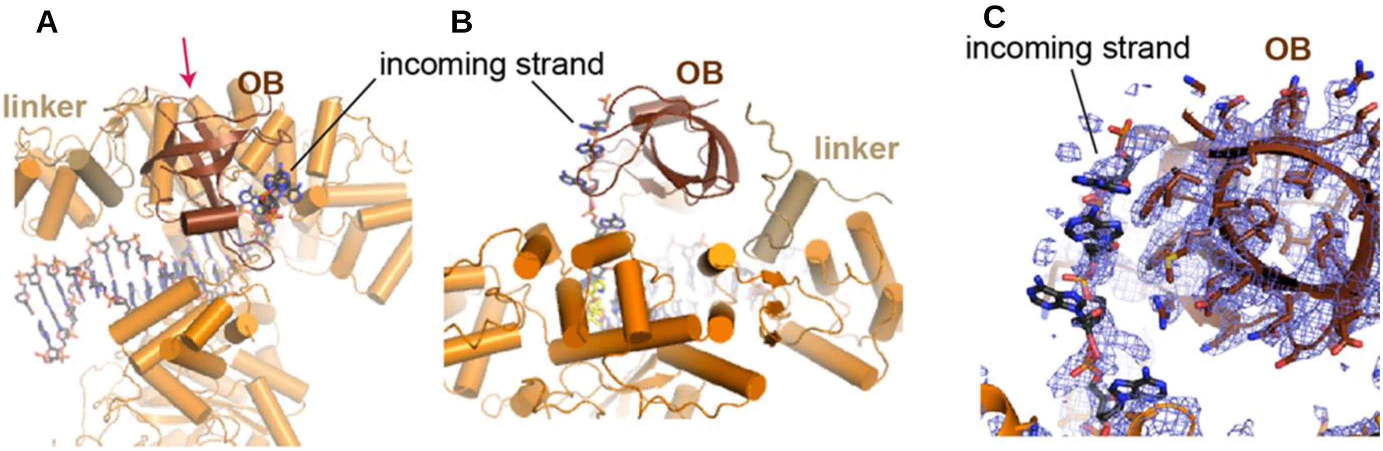


**Figure S5. The C-terminal tail of *Mtb* DnaE1 is located on top of the fingers domain. Related to Figure 4.** (A) Top view showing the position of the oligonucleotide/oligosaccharide binding (OB) domain (coloured in dark brown) over the polymerase active site and nargenicin (not visible). View is similar to that of main Figure 4B. (B) Close-up of the OB domain and the long linker between the fingers domain and OB domain as viewed along the red arrow in panel a. The connections between the linker and adjacent regions are flexible and not shown. (C) Close-up of the incoming template strand and the adjacent OB domain. The OB domain guides the incoming template strand into the polymerase active site but has little interaction with it. Cryo-EM map shown in blue mesh reveals poorly resolved density around the ssDNA.

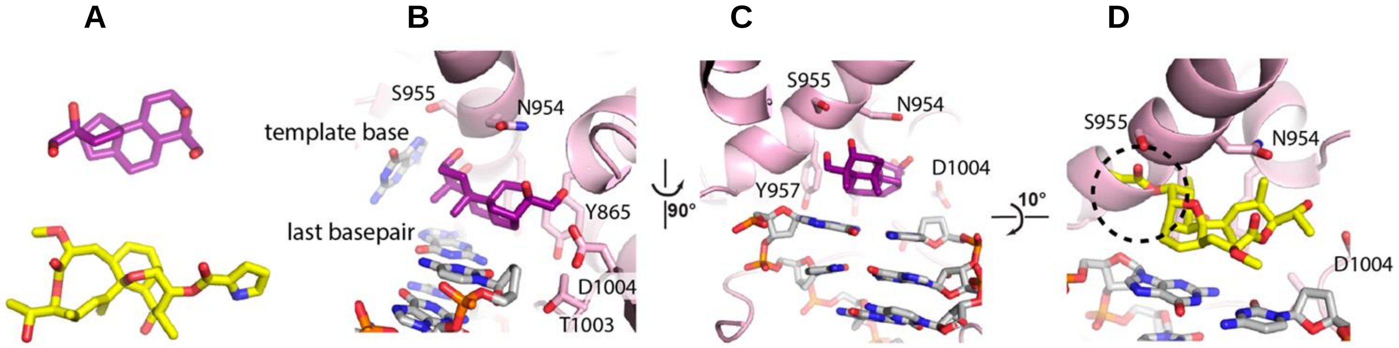


**Figure S6. Binding of aphidicolin in human Pol α. Related to Figures 4D, E and 5.** (A) Structures of aphidicolin (top, purple) and nargenicin (bottom, yellow). (B) Side view of human Pol α active site showing the position of aphidicolin wedged between the last base pair and the polymerase fingers domain. View similar to that of Figure 3D. (C) Top view of aphidicolin binding. The displaced template base is removed for clarity. View similar to that of Figure 3E. (D) Modelling of nargenicin into the human Pol α active site reveals a major clash between the pyrrole ring of nargenicin and the long helix in the polymerase, marked by a dashed circle. For the structure alignment, the DNA molecules from *Mtb* DnaE1 and human Pol α were superimposed.

**Table S1. Antimycobacterial activity of nargenicin. Related to Figure 1B and key resources table.** The activity of nargenicin against different strains of *Mtb* and *Msm* illustrates the effect of media composition on the minimum inhibitory concentration (MIC). Gene names are italicised.

| **Species** | **Strain** | **Media[a]** | **MIC (µM)** |
|---|---|---|---|
| *Mtb* | | | |
| | H37Rv | 7H9/OADC/Tw | 12.5 |
| | H37RvΔ*dnaE2* | 7H9/OADC/Tw | 6.3 |
| | H37Rv | 7H9/Glu/ADC/T | 6.3 |
| | H37Rv | GAST-Fe/Tw | 1.6 |
| | H37Rv | GAST/Tw | 1.6 |
| | H37Rv | 7H9/OADC/Tx | > 100 |
| | H37Rv | 7H9/Glu/CAS/Tx | > 100 |
| | CDC1551 | 7H9/OADC/Tw | 31 |
| | HN878 | 7H9/OADC/Tw | 31 |
| | K04b00DS | 7H9/OADC/Tw | 31 |
| | NIH_KA31b00DS | 7H9/OADC/Tw | 31 |
| | K14b00DS | 7H9/OADC/Tw | 63 |
| | K29b00MR | 7H9/OADC/Tw | 63 |
| | K33b00MR | 7H9/OADC/Tw | 31 |
| | K37b00XR | 7H9/OADC/Tw | 31 |
| | K32b00MR | 7H9/OADC/Tw | 31 |
| *Msm* | | | |
| | mc$^2$155 | 7H9/OADC/Tw | 50 |
| | mc$^2$155Δ*recA* | 7H9/OADC/Tw | 50 |
| | mc$^2$155Δ*dnaE2* | 7H9/OADC/Tw | 50 |

[a]7H9, Middlebrook 7H9 media; GAST/(Fe), glycerol alanine salts (with iron); Glu, glucose; (O)ADC, (oleic acid)-albumin-dextrose-catalase; Tw, Tween, 80; Tx, Tyloxapol.

**Table S2**. **Genes that are commonly differentially regulated *Mtb* in response to treatment with nargenicin, mitomycin C or levofloxacin, as determined by DNA microarray analyses. Related to Figure S2A and S2B.** Gene lists compared represent the top 300 genes (ranked by the average Log2 fold-change in expression data from two biological repeats. Gene names are italicised.

**Genes commonly upregulated by NRG, MMC and LEVO**

| Locus | Name | Description | Function |
|---|---|---|---|
| Rv0054 | *ssb* | Single-stranded DNA-binding protein | Essential for replication of the chromosome and involved in DNA recombination and repair |
| Rv0055 | *rpsR1* | 30S ribosomal protein S18 1 | Implicated in aminoacyl-transfer RNA binding. It appears to be situated at the decoding site of messenger RNA |
| Rv0056 | *rplI* | 50S ribosomal protein L9 | Binds to the 23S rRNA |
| Rv0058 | *dnaB* | Replicative DNA helicase | Participates in initiation and elongation during chromosome replication; it exhibits DNA-dependent ATPase activity |
| Rv0181c | Rv0181c | Putative quercetin 2,3-dioxygenase | Function unknown |
| Rv0336 | Rv0336 | Conserved 13E12 repeat family protein | Function unknown |
| Rv0427c | *xthA* | Probable exodeoxyribonuclease III protein | Involved in base excision repair |
| Rv0515 | Rv0515 | Conserved 13E12 repeat family protein | Function unknown |
| Rv0605 | Rv0605 | Possible resolvase | Prevents the cointegration of foreign DNA before integration into the chromosome |
| Rv0606 | Rv0606 | Possible transposase | Thought to be required for the transposition of the insertion element IS1536 |
| Rv0829 | Rv0829 | Possible transposase | Required for the transposition of the insertion element IS1605' |
| Rv0922 | Rv0922 | Possible transposase | Required for the transposition of the insertion element IS1535 |
| Rv1122 | *gnd2* | Probable gnd2,6-phosphogluconate dehydrogenase | Involved in hexose monophosphate shunt (pentose phosphate pathway) |
| Rv1316c | *ogt* | Methylated-DNA--protein-cysteine methyltransferase | Repair of alkylated guanine in DNA |
| Rv1317c | *alkA* | Probable bifunctional transcriptional activator/DNA repair enzyme | Involved in the adaptive response to alkylation damage in DNA caused by alkylating agents |
| Rv1377c | Rv1377c | Putative transferase | Function unknown; probably involved in cellular metabolism |
| Rv1633 | *uvrB* | UvrABC system protein B | Involved in nucleotide excision repair |
| Rv1833c | *dhmA2* | Haloalkane dehalogenase 2 | May act on a wide range of 1-haloalkanes, haloalcohols, haloalkenes and some haloaromatic compounds |
| Rv1957 | *secBL* | SecB-like chaperone | Function unknown |
| Rv2579 | *dhaA* | Haloalkane dehalogenase 3 | Generates a primary alcohol and halide from 1-haloalkane and H2O |
| Rv2719c | Rv2719c | Possible conserved membrane protein | Function unknown |
| Rv2736c | *recX* | Regulatory protein RecX | Binds to RecA inhibiting ATP hydrolysis and the generation of heteroduplex DNA |
| Rv2737c | *recA* | RecA | Involved in regulation of nucleotide excision repair, in genetic recombination, and in induction of the SOS response |
| Rv2789c | *fadE21* | Probable acyl-CoA dehydrogenase | Function unknown, but involved in lipid degradation |
| Rv2790c | *ltp1* | Probable lipid-transfer protein | Possibly catalyses the transfer of a great variety of lipids between membranes |
| Rv2791c | Rv2791c | Probable IS1602 transposase | Required for the transposition of the insertion element IS1602 |
| Rv2792c | Rv2792c | Possible IS1602 resolvase | Prevents the cointegration of foreign DNA before integration into the chromosome |
| Rv2977c | *thiL* | Thiamine-monophosphate kinase | Involved in thiamine biosynthesis |
| Rv2978c | Rv2978c | Probable transposase for IS1538 | Required for the transposition of the insertion element IS1538 |
| Rv2979c | Rv2979c | Probable resolvase for IS1538 | Prevents the cointegration of foreign DNA before integration into the chromosome |
| Rv3202c | *adnA* | AdnA subunit of DNA-resecting motor-nuclease | Has both ATPase and helicase activities |

| | | | |
|---|---|---|---|
| Rv3226c | Rv3226c | Putative SOS response-associated peptidase | Function unknown |
| Rv3296 | *lhr* | Probable ATP-dependent helicase, Lhr | Has both ATPase and helicase activities |
| Rv3394c | *imuB* | Cryptic Y-family DNA polymerase, ImuB | SOS-induced mutagenesis and damage tolerance |
| Rv3395c | *imuA'* | RecA-like protein, ImuA' | SOS-induced mutagenesis and damage tolerance |
| Rv3585 | *radA* | DNA repair protein RadA | Involved in genetic recombination. May play a role in the repair of endogenous alkylation damage |
| Rv3586 | *disA* | DNA integrity scanning protein DisA | Function unknown |
| Rv3827c | Rv3827c | Possible transposase within IS1537 element | Required for the transposition of the insertion sequence IS1537 |
| Rv3828c | Rv3828c | Possible resolvase within IS1537 element | Prevents the cointegration of foreign DNA before integration into the chromosome |
| Rv0057, Rv0059, Rv0060, Rv0184, Rv0185, Rv0959, Rv1277, Rv1702c, Rv1765c, Rv1907c, Rv1945, Rv2015c, Rv2024c, Rv2100, Rv2119, Rv2308, Rv2722, Rv2734, Rv3074, Rv3222c, Rv3466, Rv3642c, Rv3776 | | Uncharacterized | Unknown |

**Genes commonly downregulated by NRG, MMC and LEVO**

| Locus | Name | Description | Function |
|---|---|---|---|
| Rv1698 | *mctB* | Copper transporter MctB | Outer membrane channel |
| Rv2074 | Rv2074 | F420H(2)-dependent biliverdin reductase | May be involved in biosynthesis of pyridoxine (vitamin B6) and pyridoxal phosphate |
| Rv2145c | *wag31* | Cell wall synthesis protein Wag31 | DivIVA family protein Wag31 |
| Rv2147c | *sepF* | Cell division protein SepF | Part of the divisome complex and is recruited early to the Z-ring |
| Rv2150c | *ftsZ* | Cell division protein FtsZ | Essential cell division protein that forms a contractile ring structure (Z ring) at the future cell division site |
| Rv2450c | *rpfE* | Resuscitation-promoting factor RpfE | Factor that stimulates resuscitation of 'dormant' cells |
| Rv2707 | Rv2707 | Possible integral membrane protein | Probable conserved transmembrane alanine and leucine rich protein |
| Rv2986c | *hupB* | DNA-binding protein HU homolog | Histone-like DNA-binding protein which is capable of wrapping DNA to stabilize it |
| Rv3260c | *whiB2* | Transcriptional regulator WhiB2 | Transcriptional regulator of cell division |
| Rv3330 | *dacB1* | Penicillin-binding protein DacB | Involved in peptidoglycan synthesis (at final stages) |
| Rv1697, Rv3258c | | Uncharacterized | Function unknown |

**Table S3. Primers and other oligonucleotides used in this study. Related to STAR Methods** Gene names are italicised.

| Primer Name | Sequence (5'-3') | Application |
|---|---|---|
| RT-Mtb dnaE1_F | ACCGGACAACTACTTCCTTGA | Forward primer for qPCR analysis of *Mtb dnaE1* |
| RT-Mtb dnaE1_R | ACACACAACAAAGCCTCATGG | Reverse primer for qPCR analysis of *Mtb dnaE1* |
| RT-Mtb sigA_F | CGGTGATTTCGTCTGGGATGA | Forward primer for qPCR analysis of *Mtb sigA* |
| RT-Mtb sigA_R | TGCCGATCTGTTTGAGGTAGG | Reverse primer for qPCR analysis of *Mtb sigA* |
| RT-Msm dnaE1_F | CGTCGTCTCGTCACTGATCAA | Forward primer for qPCR analysis of *Msm dnaE1* |
| RT-Msm dnaE1_R | GATGCCCCAACGAATCGAAAG | Reverse primer for qPCR analysis of *Msm dnaE1* |
| RT-Msm dnaE2_F | CACACCGAGGACAGCTGG | Forward primer for qPCR analysis of *Msm dnaE2* |
| RT-Msm dnaE2_R | AACTGCTCGATGATCCTCAGC | Reverse primer for qPCR analysis of *Msm dnaE2* |
| RT-Msm sigA_F | ACCAAGGGCTACAAGTTCTCG | Forward primer for qPCR analysis of *Msm sigA* |
| RT-Msm sigA_R | CATCTCCTTGGCGAGCTCTTC | Reverse primer for qPCR analysis of *Msm sigA* |
| Mtb dnaE1sgRNA_3 | GAACGCGTAGTCAGCGAACG | sgRNA targeting sequence of *Mtb dnaE1* |
| Mtb dnaE1sgRNA_6 | AGCTGGCCCTCGAAG | sgRNA targeting sequence of *Mtb dnaE1* |
| Mtb dnaE1 sgRNA_11 | GCGGCCGCTTCCACA | sgRNA targeting sequence of *Mtb dnaE1* |
| Mtb dnaE1 sgRNA_13 | GGACCGCCACCTTGGCGTTGA | sgRNA targeting sequence of *Mtb dnaE1* |
